# Individual-specific recovery of the healthy human gut microbiota after antibiotics

**DOI:** 10.64898/2026.10.30.762915

**Authors:** Shaya Kahn, Jonathan Friedman, Amir Bashan

## Abstract

Following disruption by broad-spectrum antibiotics, the human gut microbiome undergoes a prolonged recovery where coarse ecological features, such as species richness and diversity, are largely restored. However, as these features are shared by most healthy individuals, it remains unclear whether, even after strong perturbation, individuals preferentially return toward their own pre-treatment composition (‘individual-specific recovery’) or instead switch to alternative equally healthy states. Resolving this question is fundamental for evaluating the long-term impacts of antibiotics, because an individual’s health is affected by their unique microbiome composition, affecting immune regulation and metabolic processing, so a state switch, even to a broadly healthy configuration, could carry unintended consequences. Here we compare each subject’s post-antibiotic endpoint with their own baseline and with the baselines of every other subject, across four cohorts (*n* = 121) whose regimens ranged from a five-day fluoroquinolone course to a four-day three-drug combination given to deplete the gut community. The recovered community was significantly closer to the subject’s own baseline than to the alternative healthy baselines in more than 90% of individuals under the mildest regimen and in two thirds of those under regimens that broadly target the gut anaerobes, remaining above chance in every cohort and the majority outcome even where most baseline taxa fell below detection during treatment. This specificity is not merely the persistence and fast regrowth of survivors: it persists among baseline taxa that remain undetected for weeks to months, and the same taxon re-establishes more readily in a host where it was previously detected than in one where it was not. Newly established taxa are consistently more functionally similar to the pre-treatment community than transient arrivals, and their identity correlates with the surviving community’s composition in a substantial fraction of subjects. State switching, in contrast, was uncommon and occurred almost exclusively among subjects with the largest reduction in detected taxa during the treatment. Together, these observations reveal the individual-specific nature of the microbiome resilience to broad-spectrum antibiotics, whereas the compositional changes are not random but rather functionally and ecologically associated with the subject’s baseline.

## Introduction

While highly effective against bacterial infections, broad-spectrum antibiotic treatment has the collateral effect of acutely disrupting the commensal gut microbial community, drastically reducing microbial richness and restructuring community composition within a matter of days. Following this perturbation, the microbiome typically undergoes a prolonged recovery process. Approximately three months post-treatment, within-individual alpha-diversity indices, such as richness, Shannon entropy, and evenness, generally return to their baseline values [1–6]. Similarly, beta-diversity measures indicate that the post-antibiotic community shifts back from a disrupted phase toward a healthy ecological region [1–6]. These macroscopic observations demonstrate the inherent resilience of the post-antibiotic microbial ecosystem and its capacity to restore broad features of a healthy state [1–3, 5–15].

Yet, a fundamental question remains unresolved: does the recovering microbiome preferentially return toward its own pre-treatment composition, hereafter ‘individual-specific recovery’, or switch to a different steady state (as illustrated in Fig. 1**a**)? Switching to a different microbial steady state, even one compatible with broad features of health, may alter host-associated metabolic, immune, or physiological functions. Inter-subject variability in gut microbial composition has been associated with emotion-related traits in non-clinical populations and with stress- and mood-related psychopathology in PTSD and depressive-disorder studies [16–22], as well as with mucosal and systemic immune regulation in healthy adults [23–25], and host metabolic phenotypes, including personalized dietary responses [26–29]. Therefore, understanding the spontaneous recovery trajectory and predicting its specific endpoint are crucial for informing clinical decisions following antibiotic treatment. Furthermore, uncovering these dynamics could guide precision interventions in cases where intentionally switching a patient to a different healthy state is desired.

**Figure 1:**
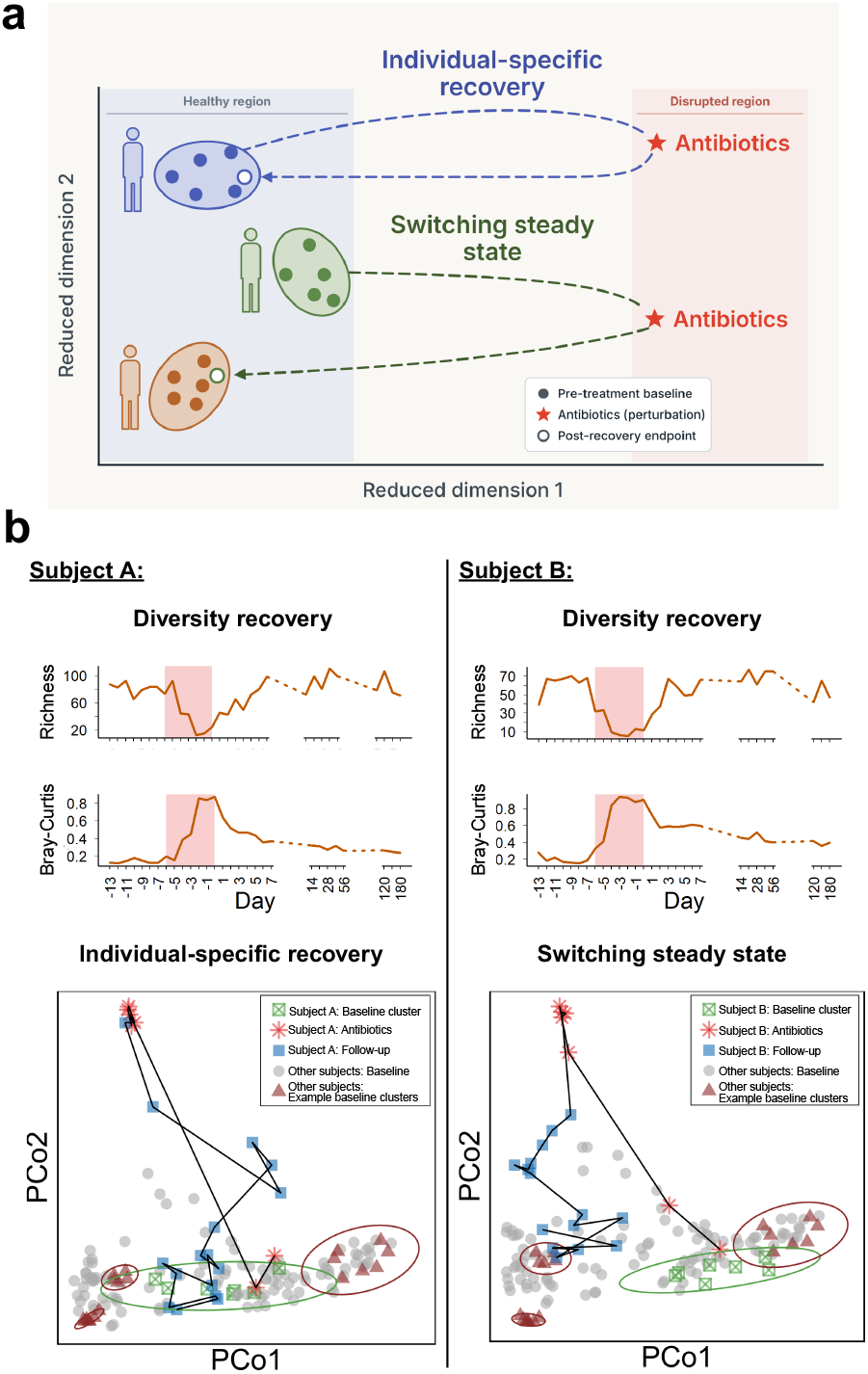
Individual-specific recovery versus switching steady state after antibiotics. **a**, Schematic of two conceptual recovery poles. In individual-specific recovery, the post-antibiotic trajectory returns toward the subject’s own pre-treatment state. In switching steady state, the trajectory moves toward a different, specific steady state or stable configuration. These poles do not define an exhaustive binary classification; intermediate or unresolved trajectories may retain weaker degrees of baseline specificity. **b**, Two example post-antibiotic trajectories from Ref. [42]. Both show restoration of broad community properties, including richness and reduced distance from the pre-treatment region, but these measures alone do not establish whether the end-point returns preferentially toward the subject’s own baseline or switches toward a different state. Subject A illustrates return toward the original baseline, whereas Subject B illustrates displacement toward a different post-treatment configuration.

While analyses based on alpha diversity or distance to baseline capture the coarse-grained transition from a disrupted community toward the healthy compositional region, they do not determine whether the endpoint preferentially approaches the subject’s own baseline or instead switches to a different steady state (Fig. 1**b**). A longitudinal case study documented a post-antibiotic shift toward a specific alternative configuration [30], whereas other studies interpreted a return to the range of natural within-subject variation as recovery toward baseline [2, 6, 31]. Distinguishing individual-specific recovery from switching steady state requires comparing each post-treatment endpoint not only with its own baseline but also with alternative healthy baselines and with the expectation under arbitrary restoration.

A tendency of a disrupted microbiome to recover toward its original state may suggest that the system retains a form of ecological memory. Such memory may arise through several complementary mechanisms. First, some species may not be completely eliminated by antibiotics but instead persist undetected at low abundance and rapidly regrow once treatment ends. Second, previously resident species may have a re-establishment advantage because they are better adapted to the host’s gastrointestinal environment and are already compatible with its immune and physiological conditions. Recovery may also be shaped by the taxa that survive antibiotic treatment. Through their ecological interactions, surviving taxa can shape the available niches, favoring the re-establishment of baseline species that are already compatible with the existing ecological network and able to persist under the prevailing competitive conditions, while limiting the establishment of less compatible taxa not detected at baseline. Finally, the functional demands of the recovering community may create additional selective pressures that favor species capable of restoring host-specific functions. Accordingly, microbiome recovery may involve not only the re-establishment of the original taxonomic composition but also the incorporation of new taxa according to their ecological compatibility and functional contributions.

In this work, we quantify the extent of individual-specific recovery by analyzing spontaneous post-antibiotic microbiome dynamics across four independent longitudinal cohorts (*n* = 121) treated with different types and doses of antibiotics. To overcome the limitations of isolated distance metrics, we employ a cross-subject comparative framework, evaluating post-treatment endpoints against both the host’s original baseline and alternative healthy configurations, defined by different subjects’ baseline states. We find that ecosystem restoration is not a random shift toward a generic healthy state: in most individuals, the recovery endpoint was significantly closer to their own pre-treatment composition than to the alternative healthy baselines. The fraction of such individuals tracked the intensity of the regimen as anticipated from the treatment parameters alone, from 93% under a five-day fluoroquinolone course that largely spares the gut anaerobes, through 81% and 61% under broad-spectrum *β*-lactams, to 33% under a four-day combination administered specifically to deplete the gut community. Individual specificity therefore weakens as the perturbation deepens, yet remains significant in the majority of subjects even when most baseline taxa fall below detection during treatment. By decoupling these recovery trajectories into distinct temporal phases, we focus on different mechanisms driving this ecological memory. We show that individual-specific recovery is initiated by the rapid rebound of suppressed taxa, but is ultimately sustained by the ecological filtering of late-returning and new species. Our results indicate that the surviving microbial community acts as an ecological filter that shapes establishment probabilities and helps define the rules governing post-antibiotic ecosystem restoration.

## Results

### Longitudinal cohorts of spontaneous post-antibiotic microbiome recovery

To determine whether post-antibiotic recovery is specifically directed toward each individual’s pre-treatment microbiome, we selected longitudinal studies that repeatedly sampled healthy subjects at baseline, during antibiotic treatment, and follow-up samples throughout the subsequent recovery period, without interventions designed to modify the recovery process. Each cohort included multiple healthy individuals whose baseline microbiomes represented distinct alternative configurations of a healthy microbial community. We analyzed data from four independent studies that tracked the spontaneous temporal dynamics of gut microbial communities: Sewunet et al. [5] (*n* = 28; baseline: day 0; antibiotics: days 2, 4, 7, and 10; follow-up: days 14, 21, 90, and 180), in which participants received either tebipenem pivoxil hydrobromide (600 mg) or amoxicillin-clavulanic acid (500/125 mg), administered orally every 8 h for 10 days (30 doses); Palleja et al. [6] (*n* = 9; baseline: day 0; antibiotics: day 4; follow-up: days 8, 42, and 180), in which participants received a combination of meropenem (500 mg), gentamicin (40 mg), and vancomycin (500 mg), administered orally as a single daily dose for 4 days; Messaoudene et al. [32] (*n* = 27; baseline: day 0; antibiotics: days 3 and 5; follow-up: days 9, 12, 16, 25, and 37), in which participants received ceftazidime-avibactam (CZA; 2 g/0.5 g every 8 h), piperacillin-tazobactam (PTZ; 4 g/0.5 g every 8 h), or ceftriaxone (CRO; 1 g every 24 h), administered intravenously for 5 days; and Yaffe et al. [13] (*n* = 57; baseline: days −63, −2, −1, and 0; antibiotics: days 1–5; follow-up: days 6, 7, 8, 10, 18, 28, and 77), in which participants received ciprofloxacin (500 mg) administered orally twice daily for 5 days. For all cohorts, microbial community composition is characterized using relative taxonomic abundance profiles (see the Methods section for details).

### Quantifying individual-specific recovery toward the pre-antibiotic microbiome

To quantify the extent to which each post-antibiotic microbial community preferentially returns toward its own pre-treatment state, we evaluated each individual using both the longitudinal and cross-sectional dimensions of the datasets. For every subject, we treated the latest available follow-up sample as the recovery endpoint and compared it with the subject’s own baseline and with a set of alternative baselines (Fig. 2**a**; see ‘surrogate data analysis’ in Methods). The alternative baselines comprise all other baseline samples available in the same study, and are not restricted to the subjects analyzed longitudinally: baseline samples of participants excluded from the longitudinal analysis, for example because their time series was incomplete, and of participants assigned to other treatment groups, were retained as additional reference healthy configurations. The number of available reference baselines, *M*, therefore equals or exceeds the number of analyzed subjects in each cohort (see Methods).

**Figure 2:**
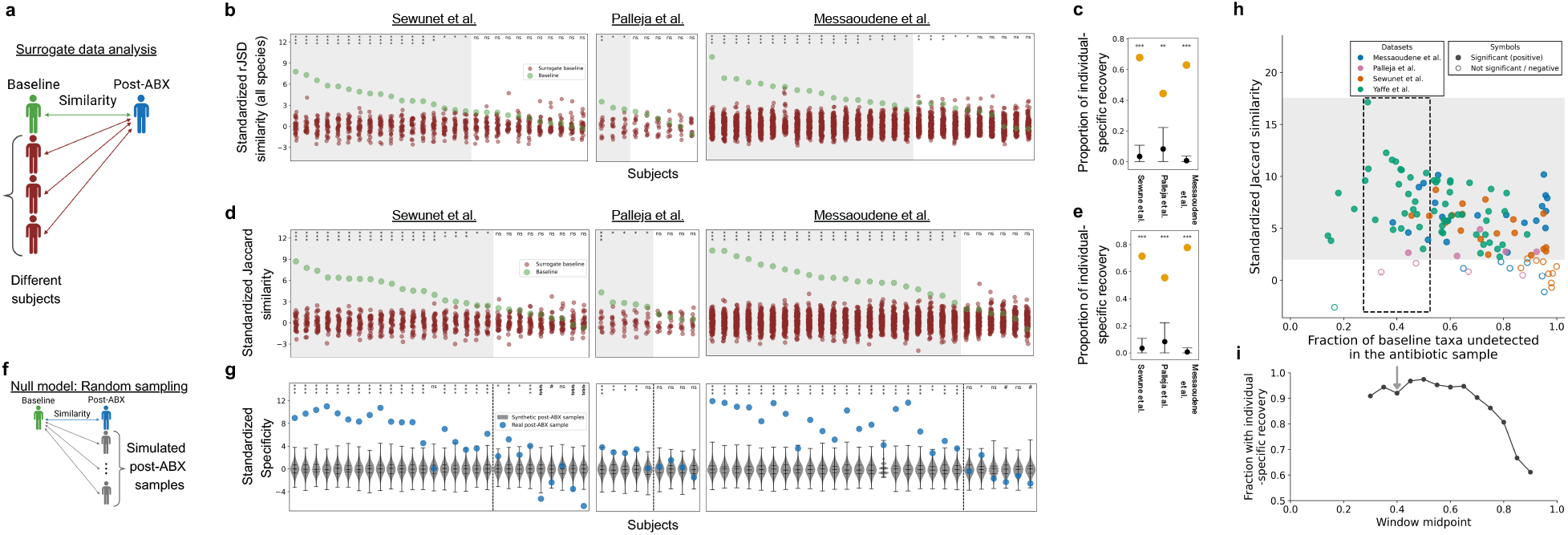
Individual-specific recovery of post-antibiotic human microbiome. **a**, Schematic demonstration of the ‘surrogate data analysis’ (see Methods). Individual-specific recovery for a subject is determined by ranking the similarity of their post-antibiotic state to their own baseline, relative to its similarity to other individuals’ baselines. **b**, Each subject’s standardized ‘rJSD similarity’ (see Methods) was calculated (large green circles). This standardization involves subtracting the mean and dividing by the standard deviation of the corresponding surrogate values from the raw rJSD similarity. The small brown circles represent the surrogate standardized rJSD similarity. The symbols above each strip plot indicate significance based on the z-score of the observed value relative to its surrogate distribution, using two-sided p-values adjusted with the Benjamini-Hochberg procedure (* * * *<* 0.001, ** *<* 0.01, * *<* 0.05, and *ns* ≥ 0.05). The significance criterion was met by 17 of 28 subjects (61%; left panel, data from Ref. [5]), 3 of 9 subjects (33%; middle panel, data from Ref. [6]), and 22 of 27 subjects (81%; right panel, data from Ref. [32]). The stricter dual criterion, which additionally requires the subject’s own baseline to rank first among all alternative baselines (grey background), was met by the same subjects in the first two cohorts and by 17 of 27 subjects (63%) in the third. The corresponding analysis for the ciprofloxacin-treated cohort of Ref. [13] (*n* = 57) is shown in Supplementary Fig. 2a. **c**, We tested whether the number of observed *rank* = 1 cases in **b** could arise by chance using a binomial test (see Methods). Yellow dots indicate the observed proportion of *rank* = 1 cases, and black dots indicate the expected proportion under the null hypothesis; error bars denote the central 90% binomial interval (see Methods). All datasets provided strong evidence against the null hypothesis (*p* = 0.004 for Ref. [6] and *p <* 10^−16^ for Refs. [5, 32]). **d**, Similarly to **b**, but using the Jaccard similarity, and excluding the survived species (see main text and Methods). As in **b**, the symbols above each strip plot indicate the significance criterion for this measure, and subjects whose own baseline additionally ranked first among all alternative baselines are shown on a grey background. The dual criterion was met by 19 of 28 subjects (68%; left panel, data from Ref. [5]), 5 of 9 subjects (56%; middle panel, data from Ref. [6]), and 21 of 27 subjects (78%; right panel, data from Ref. [32]). The corresponding analysis for the ciprofloxacin-treated cohort of Ref. [13] (*n* = 57) is shown in Supplementary Fig. 2b. **e**, As in **c**, we applied a binomial test to the results shown in **d**. All datasets provided strong evidence against the null hypothesis (*p <* 3 × 10^−4^ for Ref. [6] and *p <* 10^−16^ for Refs. [5,32]). **f**, Schematic demonstration of the ‘null model’ analysis. The ‘specificity’ between post-antibiotics and the baseline of the same subject (the fraction of species in the post-ABX that were in the original baseline, see Methods), is compared to *K* = 10^4^ values calculated between simulated samples generated from the cohort-wide baseline detection-frequency distribution under the specified statistical null model (see procedure in Methods). **g**, Each subject’s standardized specificity was calculated (blue circles) by subtracting the mean and dividing by the standard deviation of the corresponding null distribution from the raw specificity. grey violin plots represent the standardized null distribution, black marks show the median, min, max, and the 25-75% quantiles. Subjects are ordered according to their sequence in **d**. A dashed line distinguishes subjects with a statistically significant *rank* = 1 from the others. The symbols above each strip plot are defined similarly to **b** and **d**, where the # indicates cases where significance arises from the lower side of the null distribution. **h**, Standardized Jaccard similarity versus the fraction of baseline taxa undetected in the antibiotic sample. Each point represents one participant, with colors indicating datasets. Filled and open circles indicate subjects that did and did not meet the significance criterion, respectively. **i**, Fraction of subjects exhibiting individual-specific recovery across sliding windows of antibiotic disruption. Points indicate the fraction meeting the significance criterion within each window, plotted at its midpoint. Windows have a width of 0.20 and advance in steps of 0.05 along the fraction of baseline taxa undetected in the antibiotic sample. Subjects are pooled across the four cohorts.

We begin with the most basic form of this analysis and extend it in the sections that follow. Community similarity is quantified using ‘rJSD similarity’, defined as one minus the root Jensen-Shannon divergence between two microbial samples, with higher values indicating more similar relative-abundance profiles, and all detected taxa are included. Individual-specific recovery is then assessed by the significance criterion alone: a subject is classified as showing significant individual-specific recovery when its recovery endpoint is significantly more similar to its own baseline than to the alternative baselines, according to the surrogate-data analysis. This criterion is based on the continuous standardized similarity score and does not require the subject’s own baseline to be the single most similar reference, so it depends only weakly on the number of available reference baselines (Supplementary Fig. 1; see Supplementary Information). It therefore serves as our point of departure. We subsequently build on it in two directions: by adding the stricter ‘dual criterion’, which additionally requires the subject’s own baseline to rank first among all alternative baselines, and by repeating the surrogate-data analysis with alternative similarity measures and with restricted subsets of taxa.

The four cohorts differ in the expected impact of their regimens on the gut community, which can be anticipated from the treatment protocol alone. Ciprofloxacin, administered to the Yaffe et al. cohort (Ref. [13]; *n* = 57), is a fluoroquinolone that is well absorbed systemically and has limited activity against the obligate anaerobes constituting most of the gut community. The other three cohorts received agents with substantial anti-anaerobe activity that reach the colon: broad-spectrum *β*-lactams, given intravenously at therapeutic doses to the Messaoudene et al. cohort (Ref. [32]; *n* = 27) and orally over a ten-day course to the Sewunet et al. cohort (Ref. [5]; *n* = 28), and, in the Palleja et al. cohort (Ref. [6]; *n* = 9), an oral combination of three poorly absorbed bactericidal agents covering Gram-positives, Gram-negatives, and anaerobes. The fraction of subjects meeting the significance criterion followed this grouping (Fig. 2**b**). It was highest in the Yaffe et al. cohort, 93% (53 of 57; Supplementary Fig. 2**a**), the only cohort whose regimen largely spares the anaerobic core. It was intermediate in the two cohorts treated with broad-spectrum *β*-lactams, 81% (22 of 27) and 61% (17 of 28) for Messaoudene et al. and Sewunet et al. respectively. It was lowest in the Palleja et al. cohort, 33% (3 of 9), in which the combination was administered specifically to deplete the gut community; given the size of this cohort this fraction should be interpreted with caution, although it remains higher than expected by chance (see below). Thus individual-specific recovery was detected in every cohort, but its prevalence decreased as the regimen was expected to affect a larger part of the resident anaerobic community.

Applying the stricter ‘dual criterion’, which additionally requires the subject’s own baseline to rank first among all alternative baselines, left this pattern unchanged (Fig. 2**b**). Under this criterion, individual-specific recovery was observed in 89% (51 of 57) of subjects in the Yaffe et al. cohort (Supplementary Fig. 2**a**), 63% (17 of 27) in the Messaoudene et al. cohort, 61% (17 of 28) in the Sewunet et al. cohort, and 33% (3 of 9) in the Palleja et al. cohort; the corresponding unstandardized analysis is shown in Supplementary Fig. 3. The additional rank requirement reduced the fractions in the Yaffe et al. and Messaoudene et al. cohorts and left the other two unchanged, without altering the ordering across cohorts. In each of these three cohorts, the number of subjects meeting this criterion exceeded the chance expectation under a binomial test with success probability 1*/M*, where *M* is the number of reference baseline states (Fig. 2**c**; *p* = 0.004 for Ref. [6] and *p <* 10^−16^ for Refs. [5, 32]; see Methods). For the Messaoudene et al. cohort (Ref. [32]; *n* = 27), a consistent result was obtained when similarity accounted for phylogenetic relatedness among taxa, using unweighted UniFrac similarity (74% of subjects met the dual criterion; Supplementary Fig. 4). The individual-specific signal depends, however, on taxonomic resolution: in the Sewunet et al. and Messaoudene et al. cohorts it was evident at the OTU and ASV levels, respectively, but was masked when taxa were aggregated at the genus level (Supplementary Fig. 5), indicating that individual specificity resides primarily in the composition of taxa within genera. This result provides our primary evidence that post-antibiotic recovery is preferentially directed toward each individual’s pre-treatment microbial configuration. Having established this overall pattern, we next examined its scope and potential ecological basis by asking which groups of taxa contribute to it and whether individual specificity persists after specific components of the recovering community are excluded.

**Figure 3:**
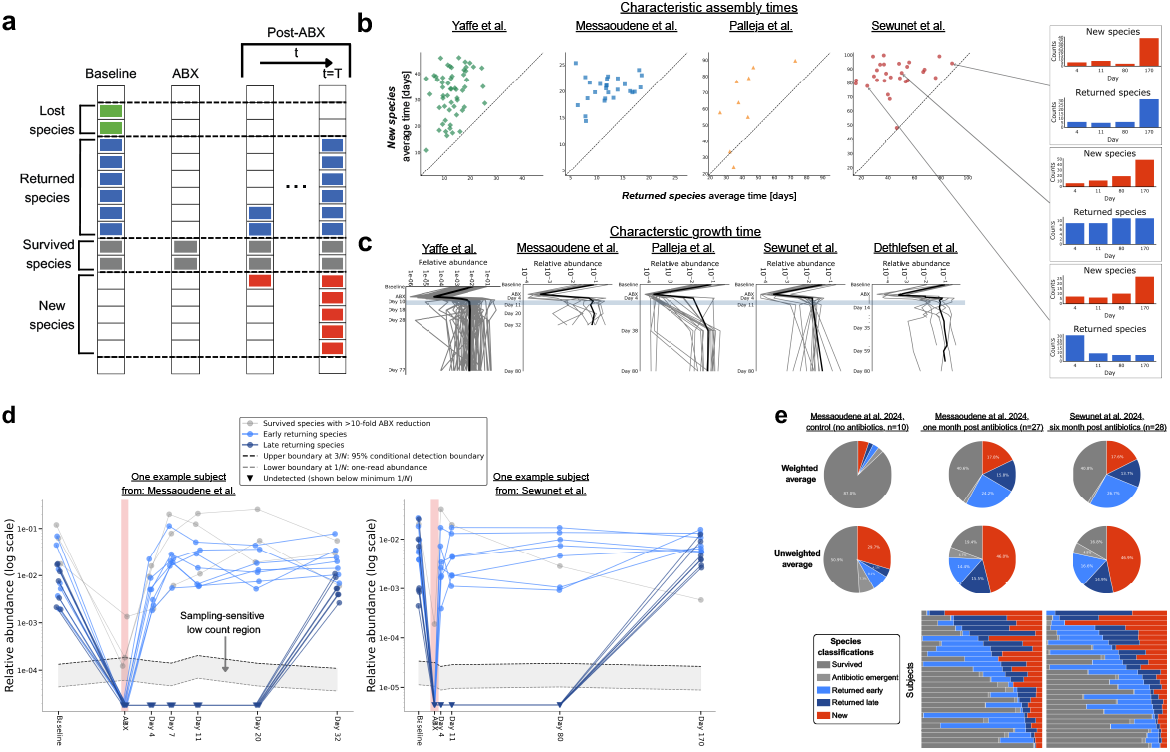
Characteristic time scales of species growth and assembly post antibiotics. **a**, Schematic of the four species categories, defined for each individual by the taxon’s longitudinal detection pattern following antibiotic perturbation (each row is a different taxon): ‘lost species’, detected in the baseline but not after antibiotic treatment (green); ‘returned species’, baseline taxa undetected during treatment but re-detected during follow-up (blue); ‘survived species’, detected in the baseline, during the antibiotic period, and at follow-up (grey); and ‘new species’, detected exclusively in follow-up samples (red). The panel is illustrative; precise operational definitions are given in the Methods. All categories refer to observed detection patterns at the applied sequencing depth. **b**, Average appearance time of the new species (*t*_*colon*_(*days*)) versus the returned species (*t*_*return*_(*days*)) associated with individual subject (see Methods). Dashed line is the equality line. The temporal emergence of new species (red bars) of three example subjects demonstrate consistent temporal dynamics across subjects, while the returned species (blue bars) dynamics is highly heterogeneous. Characteristic time scales for the appearance of new species vary by study and are affected by the duration of the observation period, as illustrated in Refs. [5, 6, 13, 32]. **c**, Recovery dynamics in logarithmic scale of the ‘suppressed species’, individual surviving species that experienced at least a 100-fold reduction in relative abundance during antibiotic treatment (see Methods, grey lines; the wide black line represents the average over the individual surviving species). The horizontal blue shaded line indicates the approximate characteristic time of abundance recovery, which is about ∼ 7 − 10 days across all five datasets (Refs [2, 5, 6, 13, 32]). **d**, Relative abundances are shown from baseline through the final antibiotic-treatment sample (ABX; pink shading) and follow-up. Early-returning species (light blue) reappeared from the first post-ABX sample onward, whereas late-returning species (dark blue) were first redetected at the final follow-up. Grey trajectories denote species detected throughout that were strongly reduced during ABX. Up to the seven most abundant species per category at the final follow-up are shown. Circles indicate detections and downward triangles non-detections. Dashed grey and black lines mark 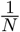 and the 95% conditional detection boundary 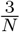 ; the shaded region between them indicates sampling-sensitive abundances. **e**, Categorization of species in a recovered microbiome. Top row: Weighted averages based on species relative abundance for antibiotic-free controls (left) and samples collected one month (center) and six months (right) after antibiotic treatment. Middle row: Pie charts showing the corresponding mean unweighted proportions of observed species across categories. Both post-treatment time points display highly similar profiles, distinct from those of untreated samples. Bottom: Stacked bar charts showing the abundance-weighted species-category distributions for individual subjects from Ref. [32] (center) and Ref. [5] (right). On average, approximately 30% of species in the recovered microbiome correspond to returned species, accounting for about 40% of total abundance; roughly half reappear within 10 days post-antibiotics (early-returning), while the remainder

**Figure 4:**
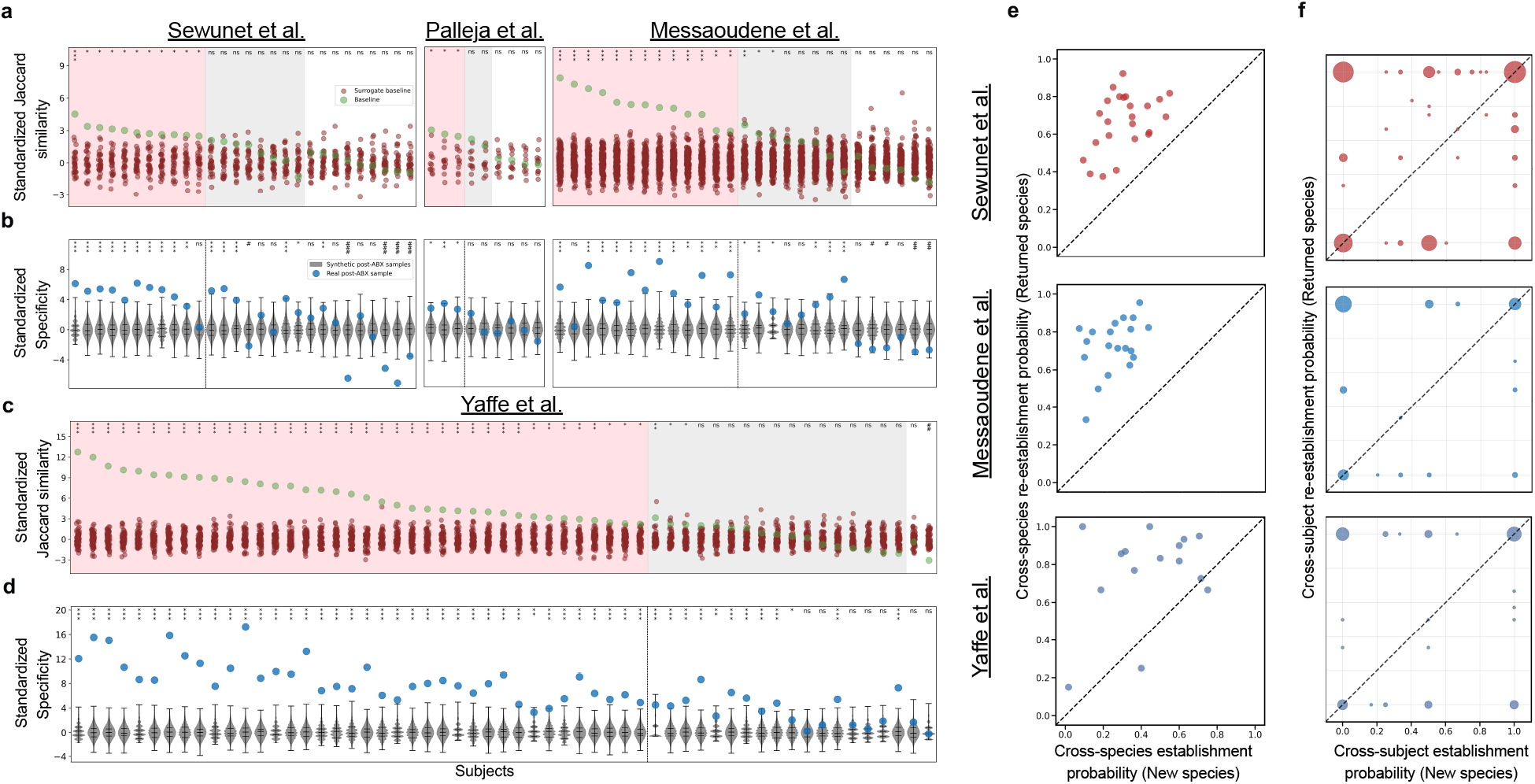
Individual-specific recovery of late-returning species in the gut microbiome post antibiotics. **a**, Similarly to Fig.2**b**, focusing only on the subsets of late-returning and new species (see Methods). Pink background represent subjects with post-antibiotics similarity with respect to their original baseline is higher than all other subjects (*rank* = 1). Grey and pink backgrounds represent subjects with *rank* = 1 when early-returning species are included or excluded, respectively. We employed a similar binomial test as depicted in Fig.2b, resulting in *p <* 0.005 for Ref. [6] and *p <* 10^−10^ for Refs. [5, 32]. **b**, As in Fig.2**g**, but with the null model applied to the reduced community of late-returning and new species (see Methods). Blue circles show each subject’s standardized specificity, obtained by subtracting the mean and dividing by the standard deviation of the corresponding null distribution; grey violin plots show the standardized null distribution, with black marks indicating the median, minimum, maximum, and 25–75% quantiles. Subjects are ordered as in **a**, and the dashed line separates subjects that met *rank* = 1 in the corresponding surrogate data analysis (**a**) from those that did not. Significance symbols above each strip plot follow Fig.2**g**; # marks subjects whose standardized specificity is significantly below the null distribution (the lower tail analysed in the main text). **c**, Similar to **a**, but for Ref. [13]. **d**, Similar to **b**, but for Ref. [13]. **e**, The panels depict the likelihood of new species and returned species observed in the penultimate sample during the follow-up period to either establish or re-establish, indicating their presence in the final sample. This is a ‘cross-species’ analysis, where new species and returned species are treated as groups, and probabilities for each subject are estimated based on group statistics. The findings are presented for subjects from Ref. [5] (top panel), Ref. [32] (middle panel), and Ref. [13] (bottom panel). Cases where probabilities were estimated based on five or fewer species were excluded. **f**, The panels are related to specific species classified as either new species or returned species that detected first at the penultimate follow-up sample, for different subjects, with each category being represented at least once. The x-axis indicates the per-case empirical probability estimate for these species when classified as new species, while the y-axis represents the same probability estimate for when the species are classified as returned species. The size of the circles is proportional in area to the number of species that occupy the same position in the figure. It is evident that the cross-subject re-establishment probabilities are generally higher than the cross-subject establishment probabilities in all datasets. A permutation test was employed to reject the null hypothesis that such a difference could occur by chance (see Methods). The results yielded *p <* 0.05 for all datasets.

**Figure 5:**
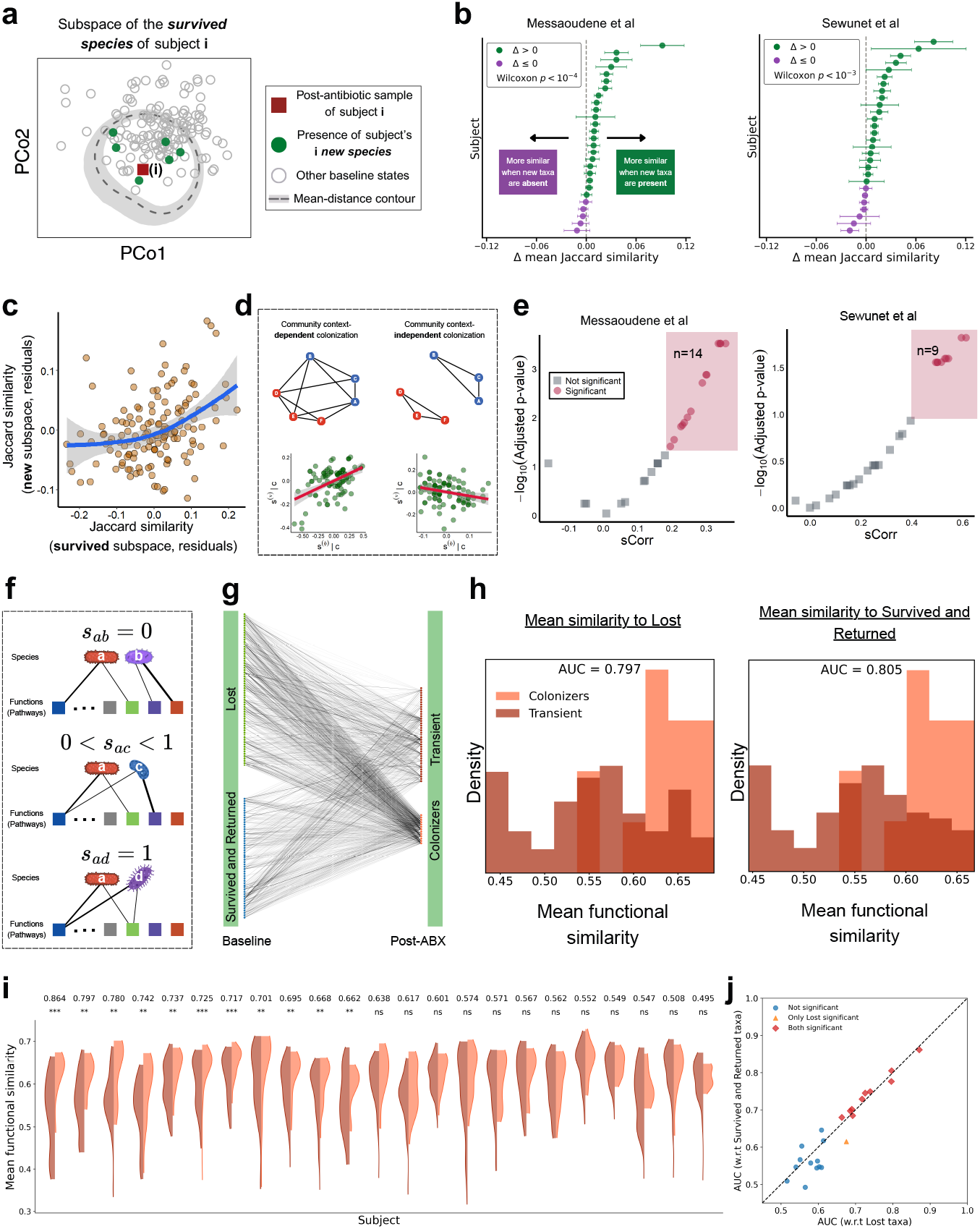
Individual-specific association between post-antibiotic new species and the baseline species. **a** PCoA based on the Jaccard distance between baseline microbiomes of different subjects (circles), calculated within the subspace of the survived species in ‘subject *i*’ (red square). The dashed curve shows the mean distance of the other subjects (grey circles) from subject *i* as a function of direction in the PCoA plane (circular kernel smoothing, *κ* = 6), and the shaded band marks its 95% bootstrap range over subjects. Green circles show the distribution of one of the new species of subject *i* across the baseline samples of other subjects; they lie mostly within the dashed curve, i.e. closer to subject *i* than the average subject in the same direction, illustrating its markedly higher prevalence among subjects that share many of the same surviving species. Data from Ref. [32]. **b**, Results are shown for Messaoudene et al. (left) and Sewunet et al. (right). For each subject and each newly detected taxon, baseline samples from other subjects were divided according to whether that taxon was present or absent. Jaccard similarity between the focal subject’s final post-antibiotic sample and each reference baseline was calculated using the subject-specific comparison set of survived, and returned taxa, excluding newly detected taxa. Reference baselines from the two groups were matched by total taxon richness, allowing a maximum difference of five taxa. For each subject, Δ is the difference between the mean similarity to baselines in which the taxon was present and the mean similarity to baselines in which it was absent, averaged with equal weighting across eligible newly detected taxa. Each point represents one subject, ordered by Δ (green: Δ *>* 0; purple: Δ ≤ 0); error bars show the standard error of the mean across that subject’s eligible taxa, and the dashed line marks Δ = 0 (no difference). The distribution of subject Δ values was tested against zero separately within each dataset using a two-sided Wilcoxon signed-rank test; the resulting *p*-values are shown in each panel. **c**, The Jaccard similarity between the post-antibiotic state of subject *i* and baseline states of other subjects is calculated in the subspace of the survived species (x-axis) and in the subspace of the new species (y-axis). A LOWESS curve is fitted. **d**, Community context-dependent colonization is illustrated in the upper panels. Species A-C form the initial community, while D-F are candidate colonizers. Interactions between the two groups induce history-dependent colonization via historical contingency. Removing these interactions renders colonization context-independent, with colonizers no longer determined by the initial community. The bottom panels show results of the historical-contingency simulation (Methods; left: *r* = 0.49, *p <* 10^−6^). When interactions are removed (right), no significant positive correlation is observed. **e**, Volcano plots of the *sCorr* values (see Methods) and their adjusted p-values. Data from Ref. [32] left panel, and [5], right panel. **f**, Functional similarity between species is defined using a bipartite species-function network, with weighted edges representing species-function associations (see Methods). In the upper panel, species *a* and *b* share no functions, yielding *s*_*ab*_ = 0. In the middle panel, species *a* and *c* share a subset of functions, yielding 0 *< s*_*ac*_ *<* 1. In the lower panel, species *a* and *d* share identical function sets with identical weights, yielding *s*_*ad*_ = 1. **g**, A network-based representation illustrates the functional similarities calculated between a subset of the ‘transient new species’ (present only in one post-ABX sample, see Methods) and the baseline species, which are divided into lost species and survived species, combined with returned species. A similar procedure is applied to the subset of ‘colonized new species’ (present in the last two post-ABX samples). The network pertains to an example subject from Ref. [32]. **h**, For the same subject from **g**, the distributions of the mean functional similarity for the colonizers and the transient w.r.t the lost species (upper panel) and w.r.t the returned and survived species (bottom panel) are shown, along with the corresponding AUC values. **i**, Each split violin represents one subject and shows the distribution of mean functional similarities for colonizers (orange, right half) and transient taxa (dark red, left half). Violin widths indicate the estimated density, with each half scaled to the same maximum width. Subjects are ordered from left to right by decreasing AUC, shown above each violin. Within-subject differences were assessed using two-sided Mann-Whitney *U* tests, with Benjamini-Hochberg correction across subjects. In this analysis, subjects with any subset of species containing fewer than 8 species were excluded. Significance symbols denote adjusted *p*-values: ^*^*p*_adj_ *<* 0.05, ^**^*p*_adj_ *<* 0.01, ^***^*p*_adj_ *<* 0.001; ns, *p*_adj_ ≥ 0.05. **j**, The AUC and the corresponding adjusted p-values for the two-sided Mann-Whitney U test were calculated for the mean functional similarity distributions of subjects from Ref. [32], subjects with any subset of species containing fewer than 8 species were excluded.

Part of the observed recovery may be attributable to the ‘survived species,’ defined as taxa detected at baseline, during antibiotic treatment, and at follow-up. To determine whether individual-specific recovery also extends beyond these persistent taxa, we repeated the analysis after excluding the survived species and retaining only the remaining taxa. Because microbiome data are compositional, removing taxa and renormalizing the retained abundances can alter sample-to-sample relationships, making conventional relative-abundance metrics sensitive to the selected subcomposition [33, 34]. We therefore converted the retained profiles to presence-absence data and quantified their similarity using the Jaccard index, a standard measure of community resemblance based on the numbers of shared and unshared taxa [35]. Because Jaccard similarity does not use abundance magnitudes, it avoids the additional rescaling introduced by renormalizing the retained subcommunity and directly measures taxonomic overlap within the operationally defined non-persistent-taxon pool. Using this presence-absence framework, individual-specific recovery met the dual criterion in 68% (19 of 28) of subjects in the Sewunet et al. cohort (Ref. [5]), 56% (5 of 9) in the Palleja et al. cohort (Ref. [6]), and 78% (21 of 27) in the Messaoudene et al. cohort (Ref. [32]) (Fig. 2**d**); the equivalent results for Ref. [13] are shown in Supplementary Fig. 2**b**, with a higher fraction of 96% (55 of 57). As above, a binomial test confirmed that these fractions exceeded the chance expectation in all three cohorts (Fig. 2**e**; *p <* 3 × 10^−4^ for Ref. [6] and *p <* 10^−16^ for Refs. [5, 32]). Nevertheless, presence-absence analysis has its own limitations: taxon detection depends on sequencing depth and the chosen detection threshold, particularly for low-abundance taxa, and binarization disregards changes in the abundances of taxa detected in both samples [36]. The Jaccard and rJSD analyses therefore provide complementary perspectives: rJSD characterizes recovery in the relative-abundance structure of the complete community, whereas Jaccard evaluates individual-specific recovery in the taxonomic membership of species not detected continuously through the perturbation. To examine whether recovery in species assemblage is nevertheless related to recovery in abundance structure, we compared presence-absence similarity with the abundance-based similarity of the species shared between baseline and post-antibiotic samples. Across the four cohorts, greater Jaccard similarity was generally associated with greater abundance similarity among the shared taxa (Supplementary Fig. 6; see Supplementary Information). Thus, although the two measures capture distinct aspects of recovery, recovery of taxonomic membership tends to coincide with recovery of the relative-abundance structure of the re-established community.

These differences among cohorts follow from what the surrogate data analysis measures. The analysis asks whether a recovery endpoint can be distinguished from alternative healthy baselines, which depends on how much of the subject’s own baseline community remained detectable through treatment. In the Yaffe et al. cohort, ciprofloxacin spared much of the obligate anaerobic community, so a large part of each baseline persisted into the recovery endpoint, which therefore retained many of the taxa and abundance relationships specific to that subject and was readily distinguished from the baselines of other subjects. In the three cohorts treated with agents active against the anaerobic core, a larger fraction of baseline taxa fell below detection during treatment and the endpoint had to be re-established from a smaller surviving community; with fewer subject-specific features carried through the perturbation, the endpoint can more easily resemble an alternative healthy baseline, and the measured specificity is correspondingly lower. The strongly disrupted cohorts therefore provide the more stringent test of directed recovery, whereas a high fraction of individual-specific recovery under a mild regimen is less informative, because part of the original community was never lost. Accordingly, the fraction of subjects showing individual-specific recovery should be read together with the extent of the perturbation rather than on its own, which we examine at the level of individual subjects below.

Given that similarities among subsets of individuals could potentially resemble individual-specific recovery, we investigated whether the observed pattern might instead indicate a broad host-associated subgroup structure. To evaluate this possibility, we analyzed baseline microbiome composition in relation to available host metadata. No significant separation was detected based on sex, age, or BMI, indicating that these measured factors do not account for the observed specificity (Supplementary Fig. 7; see Supplementary Information).

Another approach to quantifying individual-specific recovery is to test the null hypothesis that the taxa observed at the recovery endpoint were drawn under a null model from the cohort-wide baseline detection-frequency distribution. For each subject, we quantified the ‘specificity’ metric as the fraction of taxa in the post-antibiotic community that had also been detected in the subject’s own baseline (see Methods section) and compared this value with the distribution obtained from randomly generated communities matched in richness to the observed endpoint (Fig. 2**f** ; see ‘null model’ in Methods). Specificity was significantly greater than expected under the null model in 79% (22 of 28) of subjects in the Sewunet et al. cohort (Ref. [5]), 44% (4 of 9) in the Palleja et al. cohort (Ref. [6]), and 81% (22 of 27) in the Messaoudene et al. cohort (Ref. [32]) (Fig. 2**g**); the equivalent result for Ref. [13] is shown in Supplementary Fig. 2**c**, with a higher fraction of 98% (56 of 57). Significance was assessed from two-sided empirical p-values adjusted across the subjects of each cohort using the Benjamini–Hochberg procedure (see Methods).

The presence-absence surrogate-data analysis and the null-model analysis test distinct hypotheses but are both computed on taxonomic membership within the same non-surviving taxon subspace, so their subject-level classifications can be compared directly. Pooling the Sewunet et al., Palleja et al., and Messaoudene et al. cohorts (*n* = 64), 45 subjects met the dual criterion in the presence-absence surrogate analysis and 48 had significantly greater specificity than expected under the null model. The two classifications over-lapped significantly: 43 of the 45 subjects identified by the surrogate analysis were also identified by the null model (43 of 64 overall, 67%; Fisher’s exact test on the pooled cohorts, *p <* 10^−4^). In the Yaffe et al. cohort (Ref. [13]), where both analyses identified nearly all subjects (55 and 56 of 57, respectively), the two classifications necessarily coincided for at least 54 subjects, and a test of association is uninformative at these rates. Together, these findings indicate that post-antibiotic recovery is not explained solely by random sampling from a general reservoir of commensal taxa. Rather, it reflects subject-specific ecological memory and host- or microbiome-associated constraints that preferentially direct the recovering community toward the original baseline state.

The ordering anticipated from the treatment protocols is reproduced by the compositional shift measured in each cohort (see Supplementary Information). The mean Bray-Curtis dissimilarity between the final antibiotic sample and the subject’s own baseline was lowest in the Yaffe et al. cohort (0.647) and higher in the other three (0.810, 0.839, and 0.957 for the Sewunet et al., Messaoudene et al., and Palleja et al. cohorts, respectively), consistent with the colonic exposure expected from each regimen. To test whether individual specificity tracks disruption within cohorts as well as between them, we examined each subject’s standardized Jaccard similarity as a function of the fraction of baseline taxa undetected in the antibiotic sample (Fig. 2**h**). Standardized similarity decreased as this fraction increased, and subjects that did not meet the significance criterion became more frequent at higher levels of disruption; subjects whose own baseline did not rank first are indicated separately in the figure. Critically, this decline does not abolish individual specificity. Across sliding windows of disruption (Fig. 2**i**), the fraction of subjects meeting the significance criterion decreased with increasing depletion but remained above one half in every window, including the most disrupted, in which most baseline taxa were undetected during treatment. Individual-specific recovery therefore weakens as the perturbation deepens, yet remains the dominant outcome across the entire range of disruption observed, and is not merely a consequence of incomplete perturbation.

### Individual-specific recovery extends into the slow restoration phase

The analyses above show that the recovered microbiome preferentially resembles its own baseline, but they do not reveal which stages and ecological components of community restoration are responsible for this individual specificity. To address this, we separated taxa according to their observed persistence and return. This recovery-based classification is conceptually related to previous ecological classifications of microbial population dynamics, including the strain-level framework of Schmidt et al. [37], which distinguished resilience, colonization, and turnover following fecal microbiota transplantation. Here, we adapt this general perspective to longitudinal post-antibiotic recovery and distinguish taxa according to their temporal trajectories. We define ‘returned species’ as baseline taxa that were not detected during antibiotic treatment but were detected again during follow-up. These taxa likely comprise two populations: species that persisted below the detection limit and subsequently regrew, and species that were effectively absent from the community and later recolonized it. Here, ‘effectively absent’ refers to taxa that were strongly depleted by antibiotics and failed to rapidly re-establish detectable populations, regardless of whether a small residual population persisted below the detection limit. Although these categories cannot be assigned with certainty for every individual taxon, their collective behavior allows us to distinguish different contributions to individual-specific recovery.

Sequencing-based detection cannot establish the true absence of a species, and our classification therefore does not assume absolute presence or absence. Instead, it reflects whether a taxon participates detectably in the ecological community: a species that remains strongly depleted and fails to regrow may be considered effectively absent, even if a residual population persists below the detection limit. Thus, non-detection is inherently probabilistic, but repeated non-detection across consecutive samples increases the likelihood that a taxon was effectively absent. Our conclusions therefore rely on aggregate patterns across many taxa and subjects rather than on the classification of any single species.

To distinguish regrowth from slower community restoration, we used two reference subsets whose dynamics can be interpreted with greater confidence than those of the returned species. ‘New species’ were defined as taxa detected only during follow-up, neither at baseline nor during antibiotic treatment; these taxa likely reflect recruitment into the recovering community and provide an estimate of the time scale of *de novo* assembly, although some may have been present below the detection limit at baseline. In contrast, ‘suppressed species’ were baseline taxa detected at every sampled time point, including during antibiotic treatment, but whose relative abundance decreased by at least 100-fold during treatment. Because these taxa were never lost from detection, their trajectories isolate abundance recovery without requiring a re-establishment step. The two reference subsets exhibit qualitatively distinct temporal dynamics. The regrowth of suppressed species is characterized by a well-defined time scale of approximately 7-10 days, consistently observed across five independent longitudinal datasets with different antibiotic regimens and sampling designs, including one additional cohort analyzed here only for this purpose [2] (Fig. 3**c**). In contrast, the emergence of new species shows no comparable characteristic time scale. Instead, new taxa accumulate gradually over weeks to months, following a prolonged pattern that is expected to eventually approach the background turnover of an unperturbed microbiome (Supplementary Fig. 8; see Supplementary Information). Accordingly, their mean arrival time does not converge to a fixed value but increases with the duration of the observation window, consistent with a broad distribution of arrival times rather than a single dominant rate (Fig. 3**b**). Because this measure is inherently bounded by study duration, we compare it within studies rather than across studies.

Despite this prolonged accumulation process, the temporal emergence of new species is relatively consistent across subjects, with low inter-subject variability in mean arrival time for a given study duration (Fig. 3**b**). The dynamics of returned species, in contrast, are substantially more heterogeneous across individuals, consistent with different contributions of rapid regrowth and slower re-establishment. Nevertheless, in nearly all subjects, the mean return time of returned species is shorter than the mean arrival time of new species (Fig. 3**b**). Thus, the prolonged emergence of new species provides an approximate reference for the typical time scale of appearance of taxa not previously detected in the sampled community, some of which may originate from external sources. The generally faster return of returned species suggests that at least a fraction of these taxa may not have required reacquisition from an external source, but instead persisted at low abundance below the detection limit and subsequently regrew.

These measurements reveal a separation of time scales within the recovery process: a fast component, on the order of a week, associated with the rebound of populations that were suppressed but continuously detected, and a slower component, spanning weeks to months, associated with the return of taxa that were effectively absent from the community. Accordingly, we partitioned the returned species into ‘early-returning species’, redetected within the first ∼ 10 days after treatment, and ‘late-returning species’, redetected there-after. Early return is therefore more likely to reflect regrowth from populations that persisted below the detection limit, whereas late return is more consistent with re-establishment following effective absence. The threshold was estimated separately for each dataset from the observed regrowth time of the suppressed species (see Methods), and the conclusions remain unchanged when it is varied within the range permitted by the sampling resolution of each cohort. The concept of effective absence is illustrated in Fig. 3**d**. Taxa that return immediately after antibiotic treatment show dynamics similar to those of the suppressed species, suggesting that the distinction between these groups may largely reflect whether the taxon remained above or fell just below the detection limit during treatment. In contrast, taxa classified as late-returning remain undetected across all preceding follow-up samples before reappearing at substantially higher relative abundance. These reappearance abundances lie well above the sampling-sensitive low-count region (see Methods for the statistical procedure adapted from Ref. [38]). This pattern indicates that late-returning taxa undergo a substantial shift in their abundance distribution between the period of non-detection and their subsequent reappearance, supporting their interpretation as effectively absent from the detectable ecological community during the intervening period.

Consistent with this partition, species that were more likely to survive antibiotic treatment across subjects also tended to return earlier when classified as returned species, yielding a significant negative association between survival probability and average return time across all four datasets (Supplementary Fig. 9; see Supplementary Information). This supports the interpretation that early-returning species are enriched for taxa with a greater tendency to persist through treatment.

This partition offers a quantitative analysis of the composition of the recovery endpoint (Fig. 3**e**). In the cohorts studied by Messaoudene et al. [32] and Sewunet et al. [5], early-returning species constitute 14.4% (24.2% when weighted by abundance) and 16.6% (26.7% when weighted by abundance) of the taxa present at the endpoint, respectively. Conversely, late-returning species comprise 15.5% (15.8% when weighted by abundance) and 14.9% (13.7% when weighted by abundance). These comparable contributions suggest that late return is a significant component of post-antibiotic recovery, rather than a minor residual process.

The survived species and early-returning species together constitute approximately one third of the taxa present at the recovery endpoint but account for most of the community abundance, raising the possibility that they alone explain the individual specificity observed above. To test whether this specificity extends beyond the fast rebound phase, we repeated the surrogate data analysis after excluding both groups, retaining only late-returning species and new species, taxa that remained undetected throughout the rapid recovery phase (see Methods). Because this reduced community contains none of the taxa that persisted through treatment or regrew immediately afterwards, it depends far less on the extent of the initial depletion than the full community does, and we therefore report all four cohorts together. Individual specificity was reduced but remained substantial: in 54% of subjects (65 of 121) the reduced recovery endpoint still satisfied the dual criterion with respect to the subject’s own baseline (Fig. 4**a,c**). In each cohort this fraction was significantly greater than expected by chance according to the binomial test defined in the surrogate data analysis (see Methods; *p <* 0.005 for Ref. [6], *p <* 10^−10^ for Refs. [5, 32], and *p* = 2.92 × 10^−56^ for Ref. [13]). In a further 29% of subjects (35 of 121), the dual criterion was met in the full non-persistent community (Fig. 2**d**) but no longer once the early-appearing taxa were also removed, identifying cases in which specificity was primarily associated with the fast rebound phase.

Consistently, applying the null model to the same reduced community showed that 72% of subjects (87 of 121) retained significantly greater specificity than expected under arbitrary restoration (Fig. 4**b,d**). Thus, individual-specific recovery cannot be explained solely by the persistence and rapid regrowth of taxa that survived antibiotic treatment. A substantial component of this specificity persists into the slower restoration phase, among taxa that remained undetected beyond the characteristic time scale of rapid regrowth.

The lower tail of the same null distribution identified a small number of subjects whose own-baseline specificity in this reduced community was significantly lower than expected under arbitrary restoration (Fig. 4**b**). These cases therefore represent directed departures from the original baseline, consistent with a switch toward a specific alternative configuration rather than random drift. They were concentrated among subjects with the greatest antibiotic-associated loss of detectable baseline taxa (Supplementary Fig. 10). Because only a few subjects showed this pattern, and because failure to meet the significance criterion does not imply a lower-tail departure, we do not partition the full cohort into recoverers and switchers.

If the slow phase indeed carries individual specificity, a baseline taxon should be more likely to establish after reappearance in its original host than an equivalent taxon with no detected baseline history in that host. We tested this at the level of individual colonization events. For each subject, we identified taxa first detected in the penultimate follow-up sample and asked whether they remained detectable in the final sample, thereby defining an empirical persistence probability conditioned on arrival. Comparing returned species and new species at the same arrival time controls for their different characteristic time scales, so that any remaining difference is associated with the taxon’s baseline status rather than with when it appeared. Across subjects in the Sewunet et al., Messaoudene et al., and Yaffe et al. cohorts, the ‘cross-species reestablishment probability’ of returned species exceeded the ‘cross-species establishment probability’ of new species (Fig. 4**e**); estimates based on five or fewer taxa were excluded. Because lower-abundance taxa may be less likely to be detected, we further repeated the analysis across increasing minimum relative-abundance thresholds. The same pattern was retained across thresholds (Supplementary Fig. 11**a**; see Supplementary Information), indicating that differences in detectability are unlikely to explain the observed re-establishment advantage.

A remaining possibility is that returned species and new species are intrinsically different groups of organisms, for example because baseline taxa are generally more prevalent or better adapted to the gut. We therefore inverted the analysis and performed the comparison within taxa. We identified taxa that occurred as a returned species in at least one subject and as a new species in at least one other, allowing each taxon to serve as its own control, and compared its ‘cross-subject re-establishment probability’ with its ‘cross-subject establishment probability’ (see Methods). In all three of these datasets, re-establishment probability was consistently higher (Fig. 4**f** ; *p <* 0.05 for all datasets, permutation test; see Methods). Because taxon identity is held fixed while host identity varies, this difference cannot be attributed simply to intrinsic properties of returned versus new taxa: the same taxon is more likely to establish in a host in whose baseline it was detected than in a host to which it is new. Consistently, we found no significant difference in relative abundance between taxa that persisted and those that appeared only transiently, for either returned species or new species (Supplementary Fig. 11**b**; see Supplementary Information).

Together, these results support the existence of ‘ecological memory’ in microbiome recovery, whereby baseline-resident taxa are preferentially re-established relative to non-resident taxa beyond what can be explained by persistence through treatment, population prevalence, arrival time, or abundance-dependent detection. Importantly, this memory is not confined to taxa that survive the perturbation and rapidly regrow; it extends to species redetected weeks after treatment, on time scales comparable to those of taxa newly detected in the sampled community. Whether this preference arises from ecological filtering by the surviving community, host-specific physiological conditions, or individual differences in microbial exposure remains unresolved.

### Historical contingency as a mechanism shaping the establishment of new species

The surrogate data analysis investigates whether a subject’s post-antibiotic state more closely resembles its own baseline than the baselines of other subjects. New species are, by definition, not detected in the subject’s sampled baseline, resulting in a similarity of zero by construction. Consequently, the comparison used to define individual-specific recovery cannot be applied to these new species. To determine whether their assembly is also individual-specific, a different question must be posed.

We asked instead whether the identity of the new species is predictable from the community that survived the perturbation. Under a historical-contingency mechanism, the species able to establish in a depleted community are constrained by the species already present, through the ecological interactions between them (Fig. 5**d**, upper panels). If this holds, two subjects whose surviving communities resemble each other should tend to recruit similar new species, and the new species of a given subject should be found preferentially in the baselines of the subjects whose surviving communities resemble its own. If instead the recruitment of new species is independent of community context, their identity should carry no such association.

To test this, we developed a measure we term the ‘similarity correlation’ (*sCorr*). For a focal subject, we compare its post-antibiotic state with the baseline states of all other subjects twice: once restricted to the species that survived in the focal subject, and once restricted to its new species. This yields two similarity values per reference subject, and *sCorr* is the correlation between them across all references (Fig. 5**c**). A positive *sCorr* means that the subjects whose communities most resemble the focal subject’s survivors are also the subjects most likely to harbour its new species, which is the signature expected under community-dependent recruitment of new species. The principle is illustrated in Fig. 5**a**, which shows, for one subject, that a particular new species is concentrated among the reference subjects that share many of that subject’s surviving species. Across subjects, the final post-antibiotic community was on average more similar to baselines in which a given new species was present than to richness-matched baselines in which it was absent (Fig. 5**b**; two-sided Wilcoxon signed-rank test, *p <* 10^−4^ for Ref. [32] and *p <* 10^−3^ for Ref. [5]).

Before applying this measure to the data, we verified that it detects the mechanism it is intended to detect, using a simulated community model (generalized Lotka-Volterra; see Methods). The simulation does not represent the antibiotic treatment itself. It begins from the state the treatment leaves behind: each simulated community is assembled from a subset of a shared species pool and allowed to reach equilibrium, and this depleted equilibrium community stands for the surviving community of one subject. All species absent from a community are then supplied at a low immigration rate from the shared pool, representing the continuous exposure of the recovering gut to an external reservoir, and the community is allowed to reach a new equilibrium. The species that establish above the detection threshold in this second equilibrium are the simulated new colonizers, and the remaining parameters are given in Methods.

Applied to these simulations, *sCorr* was positive and highly significant (*r* = 0.49, *p <* 10^−6^; Fig. 5**d**, lower left panel). We then repeated the simulation with a single change: the interactions between the resident species and the species arriving from the pool were removed, while all other parameters were held fixed. The correlation disappeared (Fig. 5**d**, lower right panel). Because the species pool, the immigration rate and the detection threshold were identical in the two cases, the association cannot arise from shared exposure or from the structure of the measure itself, and is attributable to the interactions between the resident community and the incoming species. To confirm that this behaviour does not depend on the particular form of the interaction model, we repeated the analysis using a mechanistic consumer-resource model, in which competition emerges from the sharing of a finite set of resources rather than being imposed through fixed interaction coefficients. The same positive association was recovered (*r* = 0.43, *p <* 10^−3^; Supplementary Fig. 12 and Supplementary Information).

We then applied the measure to the two cohorts with sufficient sampling [5, 32], treating each subject’s final follow-up sample as the post-antibiotic state and the baselines of all other subjects as references. The surviving community of a subject was defined as the union of its survived species, the taxa that emerged during the antibiotic phase, and its early-returning species, that is, the taxa present once the fast rebound phase was complete (see Methods). A positive and significant *sCorr* was found in 42% of cases (23 of 55; Fig. 5**e**). The new species that appear in a recovering gut are therefore not drawn at random from those available to it: in a substantial fraction of subjects, their identity is associated with the particular community that survived the perturbation.

This provides a mechanism through which individual specificity can extend to taxa with no prior history in the host. The community left behind by the antibiotics is associated with what subsequently becomes established, so that the surviving taxa shape not only their own recovery but also the composition of the newly arriving ones. It also suggests why baseline taxa re-establish so readily: having previously coexisted with the survivors, they are by construction compatible with the ecological context in which they reappear.

### Functional association between new colonizers and baseline species

The previous section showed that the identity of the new species is associated with the composition of the community that survived the perturbation. A natural interpretation is that these species are selected for what they do rather than for what they are, and in particular that they restore functions carried by baseline taxa that were no longer detected after antibiotics. We therefore asked whether the new species that succeed in establishing are functionally distinguishable from those that do not.

To address this we characterised species by their functional repertoires rather than by their taxonomic identity. Following Ref. [39], we represented the relationship between species and functions (MetaCyc path-ways) as a bipartite network in which weighted edges link each species to the functions encoded in its genome. The ‘functional similarity’ between two species is then the weighted Jaccard similarity between their functional profiles, which is 0 for species sharing no functions, 1 for species with identical profiles and weights, and intermediate for partial overlap (Fig. 5**f** ; see Methods).

Within each subject, we separated the new species by their fate. ‘Colonizers’ are new species present in the last two post-antibiotic samples, and ‘transient’ are new species detected in only a single post-antibiotic sample and absent from the final one. Both groups arrived in the same gut over the same period, so they are exposed to the same regional pool and the same host; they differ in whether they established. For every new species we computed its mean functional similarity to a reference set of baseline taxa, which assigns each species a single score. Two reference sets were used: the ‘lost species’, present at baseline but detected neither during treatment nor afterwards, and the ‘survived and returned’ species, which persisted through the perturbation or reappeared after it. The first set tests whether colonizers specifically replace what was lost; the second tests whether they simply resemble the baseline community in general (Fig. 5**g, h**).

We quantified the separation between the two groups using the area under the receiver operating characteristic curve (AUC) of the mean ‘functional similarity’ (see Methods). The AUC does not depend on any particular threshold: it is equal to the probability that a randomly chosen colonizer has a higher mean functional similarity to the reference set than a randomly chosen transient species. An AUC of 0.5 therefore indicates that functional similarity carries no information about which species establish, while values above 0.5 indicate that colonizers are the more functionally similar of the two groups. Significance was assessed with a two-sided Mann-Whitney *U* test on the same scores, with p-values adjusted across subjects (see Methods). Subjects with fewer than eight species in any of the relevant subsets were excluded, leaving 22 subjects from Ref. [32].

The separation was significant in 45% of the subjects tested (10 of 22; Fig. 5**i, j**). In every subject, the AUC exceeded 0.5 for both reference sets, so wherever functional similarity was informative, it was the colonizers that were the more similar group. Crucially, the two AUC values track each other closely and fall near the identity line (Fig. 5**j**): a subject whose colonizers resemble the lost species is, to a similar degree, a subject whose colonizers resemble the survived and returned species. Only one subject was significant with respect to the lost species but not with respect to the survived and returned species, and none showed the converse. A consistent pattern was obtained using functional profiles derived from shotgun metagenomic data of Ref. [13] (Supplementary Fig. 13; see Supplementary Information).

These results do not support the hypothesis we began with. Successful colonizers are not specifically matched to the functions of the ‘lost species’, baseline taxa strongly depleted during treatment and not re-established afterward; they are simply more functionally similar to the baseline community as a whole. What predicts establishment is therefore general functional compatibility with the resident community rather than the filling of a specific functional gap. This is consistent with the community-context effect of the previous section, and it offers an interpretation of why baseline taxa re-establish so readily: having been members of that community, they are guaranteed to be compatible with it in exactly this sense.

## Discussion

Our analyses revealed three main findings. First, most individuals preferentially recovered toward their own pre-antibiotic baseline rather than toward an alternative healthy configuration, and the fraction of such individuals tracked the impact expected from the antibiotic regimen, declining as the agents administered acted more strongly on the obligate anaerobes that constitute most of the gut community. Second, individual-specific recovery is observed not only in the rapid rebound immediately after treatment but also in the slower restoration phase. Third, the identity of new colonizers was associated with the surviving community, while successful colonizers tended to be functionally similar to baseline taxa.

The persistence of individual-specific recovery during the slow restoration phase suggests that baseline-directed recovery is not driven solely by the rapid regrowth of taxa that survive antibiotic treatment. One possible explanation is that the surviving community acts as an ecological filter on subsequent establishment. Consistent with this interpretation, the identities of new species were associated with the surviving community in a substantial subset of subjects, and successful ‘colonizers’ tended to be more functionally similar to baseline taxa than transient arrivals. The same filtering process could explain the home-host advantage of returned species: taxa that previously coexisted with a host’s community may be better suited to the ecological context remaining after treatment and therefore more likely to re-establish in their original host than in another host in which they were not detected at baseline. Importantly, interpreting late-returned species as effectively absent does not require assuming their complete elimination. Even if some persisted below the detection limit, the large number of newly appearing taxa that failed to persist indicates substantial exposure to alternative taxa that could potentially establish themselves in the community during recovery (Supplementary Information, Section 8). Thus, the preferential establishment of baseline-associated taxa occurs in a setting where many alternative taxa were detected transiently but did not remain detectable, supporting a selective rather than purely exposure-limited recovery process. Residual populations may still provide an additional establishment advantage, but they are unlikely to explain the full pattern. Resident-dependent filtering may therefore provide a common mechanism linking the selective return of baseline taxa to the recruitment of new species, although our observational analyses do not establish this mechanism causally.

Switching to a different post-antibiotic configuration was uncommon among the healthy individuals and antibiotic regimens examined here. Nevertheless, a small number of subjects showed a directed departure toward a specific alternative configuration: in the null-model analysis restricted to the slow-restoration community of late-returning and newly detected taxa (Fig. 4**b**), their endpoint specificity was significantly below the random-sampling null distribution, indicating a stronger departure from baseline than expected under arbitrary restoration (Supplementary Fig. 10). These cases were concentrated among subjects with the greatest antibiotic-associated loss of detectable baseline taxa, consistent with the possibility that severe disruption weakens the ecological constraints favoring return toward the original state. Severe depletion was not sufficient to determine the outcome, however, because several subjects with comparably severe antibiotic-associated depletion nevertheless recovered toward their baselines. Switching therefore depends on factors beyond perturbation magnitude, potentially including which taxa remain detectable, host-associated filtering, residual populations below detection, and the timing and composition of subsequent microbial exposure. The lower-tail null-model result establishes a directed, non-random departure but does not identify the alternative configuration or the mechanism that produced it.

Taxa classified as ‘returned species’ remained undetected for an interval before reappearing, but the available sampling cannot determine whether they persisted below the detection limit or were reacquired from an external source. Distinguishing these possibilities is important because they imply fundamentally different explanations for what changed to enable their late return. Reacquisition would depend on renewed exposure to a regional source pool followed by successful colonization. By contrast, regrowth of populations that persist through antibiotic treatment may involve resuscitation from dormant or low-activity states [40, 41], or indicate that conditions within the gut had changed sufficiently to permit their recovery, potentially through the sequential recovery of other resident taxa, restoration of ecological interactions, and interactions with newly establishing species. Strain-resolved comparisons could help distinguish these scenarios: recovery of a lineage genetically matching the baseline population would support persistence or reacquisition of the same strain, whereas the appearance of a genetically distinct but closely related lineage would be more consistent with colonization by a taxon with similar ecological or functional properties. Denser longitudinal and strain-resolved sampling, together with measurements of potential source pools, will be required to distinguish these mechanisms and identify the triggers of late reappearance.

Several features of the study limit the scope of the conclusions, including the modest cohort sizes, ranging from 9 to 57 subjects, and the unequal follow-up periods, ranging from 37 to 180 days. The four antibiotic protocols differ in route, dose, duration, and spectrum, and we use these differences to order the cohorts by their expected impact on the gut community; this ordering, however, rests on pharmacological reasoning rather than on colonic drug concentrations measured in our own subjects, and two of the three resulting groups contain a single cohort. Within the Messaoudene et al. cohort, moreover, the three agents administered are not equivalent in this respect, since ceftriaxone reaches the colon at substantially lower active concentrations than ceftazidime-avibactam or piperacillin-tazobactam [32]. The operational categories ‘survived species’, ‘lost species’, ‘returned species’, ‘new species’, ‘suppressed species’, ‘early-returning species’, and ‘late-returning species’ depend on sampling frequency, detection limits, and taxonomic resolution, and therefore should not be interpreted as certain biological states. Although presence-absence and abundance-based analyses gave consistent evidence for baseline-directed recovery, neither resolves strain identity, and this limitation is particularly important in the marker-gene cohorts. The mechanistic analyses are also more restricted in scope than the analyses of individual-specific recovery: the similarity-correlation analysis was applied to two cohorts and the functional comparison of colonizers and transient taxa to one, with a pooled replication in a third. Moreover, the observational data cannot separate filtering by the resident community from host physiological or immune filtering, differential exposure to source pools, or combinations of these processes. The GLV analysis establishes the plausibility of interaction-mediated historical contingency but does not identify a universal interaction network or prove that this mechanism dominates in humans. Controlled perturbation experiments, including gnotobiotic models with defined resident communities and exposure pools, will be required to distinguish these mechanisms and test their causal contributions.

At the population level, approximately 73% of subjects (40 of 55) in the Sewunet et al. [5] and Messaoudene et al. [32] cohorts meet the dual criterion for individual-specific recovery when quantified by the standardized Jaccard similarity after survived species are excluded, providing a baseline expectation for recovery outcome. This expectation can be refined at the individual level: in Supplementary Fig. 14, we trained regularized linear models on these two cohorts, using each subject’s baseline taxonomic composition to predict this standardized Jaccard similarity score, and evaluated performance by cross-validation. Predicted and observed recovery scores were positively correlated in both cohorts, with Pearson correlations of *r* = 0.574 for Sewunet et al. and *r* = 0.544 for Messaoudene et al. (Supplementary Fig. 14; see Supplementary Information). These results suggest that baseline community composition contains information that can shift the expected recovery outcome for a given individual above or below the population-level average. Identifying subjects at higher risk of incomplete recovery or a potential state switch could ultimately help guide post-antibiotic monitoring or intervention. Although the clearest state-switch-like endpoints occurred only after high measured baseline-taxon depletion, substantial perturbation alone was not sufficient to determine the outcome, suggesting that useful prognostic models will likely need to combine perturbation magnitude with baseline microbiome composition and other host or environmental factors.

## Methods

### Surrogate data analysis

Consider a longitudinal experiment involving subjects, sampled at baseline, during antibiotic administration, and in the post-antibiotic phase. Additionally, *M* reference baseline samples are available. We define the set of samples for subject *ν* in the experiment as [**b**^(*ν*)^, **a**^(*ν*)^, **p**^(*ν*,1)^, …, **p**^(*ν,T*)^], where 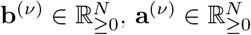, and 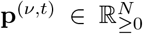 represent the baseline, the last sample of the antibiotic phase, and sample number *t* after antibiotics, relative abundance samples, respectively. To implement the ‘surrogate data analysis’ for a specific test subject *ν*, we executed the following steps: We defined the set *w* = *u* ∪ *s*, where 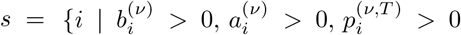, | *i* = 1, 2, …, *N* } is the set of ‘survived species’ of subject *ν*, and 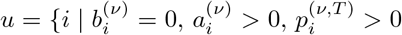, | *i* = 1, 2, …, *N* }, is the set of ‘antibiotic emergent’ species of subject *ν*. Subsequently, we computed the Jaccard similarity 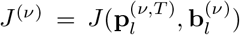, where *l* = *w*^*c*^ denotes the complement of *w*. Following this, we calculated 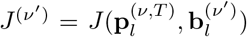 where *ν*^′^ ∈ {1, …, *M* } and *ν*^′^≠ *ν*. Finally, we determined the *rank* using the formula:

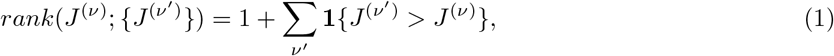

where **1**{condition} is the indicator function that equals 1 if *condition* is true and 0 otherwise. The procedure above is stated in its general form, in which a subset of taxa is excluded through the set *w*. In the main text it is applied first with an abundance-based similarity measure, the rJSD similarity (i.e., one minus the rJSD dissimilarity), computed over all detected taxa by setting *w* = ∅, and subsequently with the Jaccard similarity over the restricted taxon subsets defined above and in the sections that follow.

### Standardized similarity score and significance

For a test subject *ν*, let *S*^(*ν*)^ denote the similarity between the recovery endpoint and the subject’s own baseline (the Jaccard similarity *J*^(*ν*)^ defined above, or the rJSD similarity in the abundance-based analysis), and let 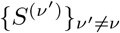 denote the corresponding similarities to the *M* − 1 alternative reference baselines. We standardized the own-baseline similarity relative to this surrogate set,

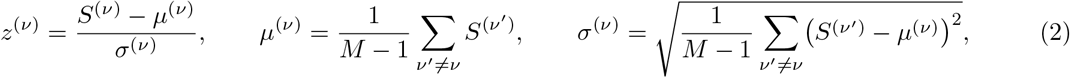

using the population standard deviation. A positive *z*^(*ν*)^ indicates that the recovery endpoint is more similar to the subject’s own baseline than to the average alternative baseline; the standardized surrogate values 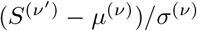 form the reference distribution shown in Fig. 2**b,d**. Treating *z*^(*ν*)^ as a standard-normal statistic, we computed a two-sided p-value

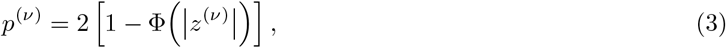

with Φ the standard normal cumulative distribution function, and adjusted the p-values across the subjects of each dataset with the Benjamini–Hochberg procedure at a false discovery rate of *α* = 0.05, obtaining adjusted values 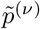. A subject was classified as showing significant individual-specific recovery when 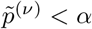 and *z*^(*ν*)^ *>* 0. The stricter dual criterion additionally requires that the subject’s own baseline rank first, *rank S*^(*ν*)^; 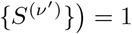.

### Binomial test for *Surrogate data analysis*

To evaluate the null hypothesis, which posits that the group-level results of the surrogate data analysis for each dataset can be obtained randomly, we employed a binomial test using the `binomtest` function from the `scipy.stats` library in Python. We defined the number of successes as the number of subjects, *N*, achieving a *rank* = 1 in the surrogate data analysis, the number of trials as the total number of subjects tested, and the success probability as 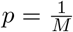, where *M* represents the number of reference baseline states (including the baseline of the tested subjects). We utilized the right-sided p-value to determine the rejection of the null hypothesis. The central 90% binomial interval for the number of successes expected under the null hypothesis, assuming a binomial distribution with parameters *N* and *p*, computed using the `binom.interval()` function from the `scipy.stats` library in Python.

### Null model: Random sampling

To implement the ‘null model’ for a specific test subject *ν*, we executed the following steps: We generated *K* simulated post-antibiotics samples 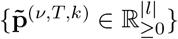 where *k* = 1, …, *K*. based on random sampling from the baseline samples of the *M* subjects, in the subspace *l*: 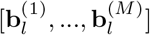, such that species are sampled based on their presence distribution, under the constraint that the number of present species in each simulated sample 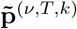 is similar to the number of present species at 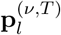. Subsequently, we calculate the ‘specificity’, 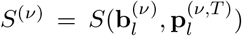. For each of the *K* simulated samples, we then compute the specificity, 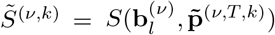. Following this, we calculate *Ŝ*^(*ν*)^, the ‘standardized specificity’ (z-score), and the ‘two-sided empirical p-value’ using the following formulas:

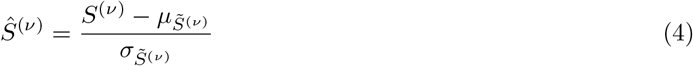

where

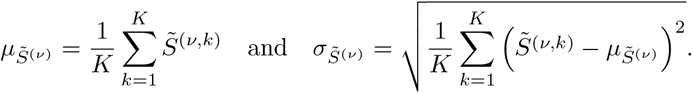

and

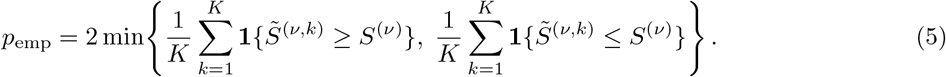

As in the surrogate-data analysis, the two-sided empirical p-values *p*_emp_ were adjusted across the subjects of each dataset with the Benjamini–Hochberg procedure at a false discovery rate of *α* = 0.05. A subject was classified as showing significantly greater-than-null specificity when the adjusted p-value was below *α* and *Ŝ*^(*ν*)^ *>* 0; the complementary significant cases with *Ŝ*^(*ν*)^ *<* 0, in which endpoint specificity is significantly *lower* than expected under random sampling, are marked with ‘#’ in Fig. 2**g** and Fig. 4**b**.

### Average appearance time

To characterize the temporal dynamics of new species, we computed the average time required for the appearance of *S* new species in each subject: 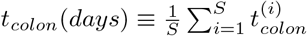, where 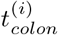 denotes the estimated appearing time of the new species *i*, i.e., the midpoint between the last sample in which it was not detected and the first sample in which it was detected. (This time scale necessarily depends on the study duration, since 1 *< t*_*colon*_ *< D*^(*T*)^, where *D*^(*T*)^ is the last day of sampling.) Similarly, we computed the characteristic time scale for returned species, *t*_*return*_(*days*).

### Identification of returned species and sampling model

To describe the effect of finite sequencing depth on species detection, we use a binomial sampling model and assume that the underlying relative abundance of a species is fixed within a given sample. Thus, this calculation considers sampling noise alone and does not account for additional temporal or ecological fluctuations in species abundance. Let *x*_*i*_ denote the underlying relative abundance of species *i* in a sample containing *N* reads. The observed number of reads assigned to that species, *n*_*i*_, is modeled as

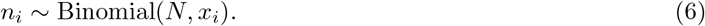

The probability of failing to detect the species is therefore *P* (*n*_*i*_ = 0 | *x*_*i*_, *N*) = (1 − *x*_*i*_)^*N*^, and the probability of detecting it at least once is *P* (*n*_*i*_ ≥ 1 | *x*_*i*_, *N*) = 1 − (1 − *x*_*i*_)^*N*^ . The relative abundance corresponding to a 95% probability of detection is

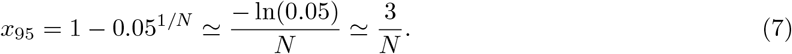

We therefore use 3*/N* as an approximate 95% conditional detection boundary. The value 1*/N* corresponds to an expected read count of one, and abundances between 1*/N* and 3*/N* are treated as sampling-sensitive. These boundaries quantify uncertainty due to finite sequencing depth under the assumption of a fixed underlying abundance and should not be interpreted as evidence of true biological absence.

### Surrogate data analysis and Null model: Late-returning species

In order to conduct the surrogate data analysis and the null model with a focus solely on the ‘late-returning species’, we define the set of returned species that reappeared after *D*^(*t*)^ days (the number of days following antibiotic treatment corresponding to index *t*) for a given test subject *ν* as 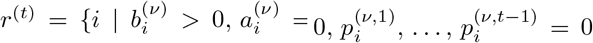, | *i* = 1, 2, …, *N*, | *t* = 1, 2, …, *T* }. The ‘early-returning species’ are characterized by the set 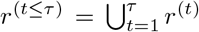, where *τ < T* . This is calculated empirically for each dataset using the estimated regrowth time of the ‘suppressed species’ (see main text), which is defined as 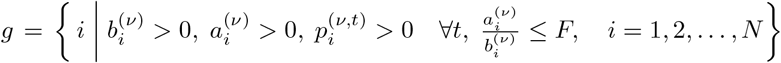, where *F* = 0.01. Similarly, we define the late-returning species as *r*^(*t>τ*)^. Subsequently, we applied the surrogate data analysis and the null model as previously described, but defined *q* = *u* ∪ *s* ∪ *r*^(*t*≤*τ*)^, and used *q* in place of *w*.

### *Cross-species* and *Cross-subject* re/establishment empirical probability analysis

To determine the ‘cross-species re-establishment empirical probability’ for a specific subject, we define the set of species 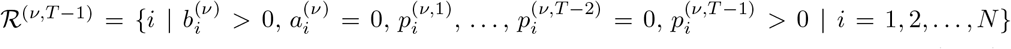. The cross-species re-establishment empirical probability is then defined as 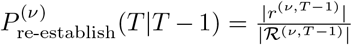, which represents the proportion of species that reappeared at timepoint *T* − 1 and are also present at timepoint *T* . The ‘cross-species establishment empirical probability’ is determined in a manner analogous to the calculation of 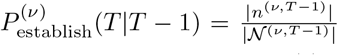, where *n*^(*ν,T* −1)^ and *N* ^(*ν,T* −1)^ are defined similarly to *r*^(*ν,T* −1)^ and ℛ^(*ν,T* −1)^, with the condition that 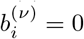 instead of 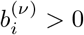 for *i* = 1, 2, …, *N* .

To determine the ‘cross-subject re-establishment empirical probability’ for a given species, we identify a specific set of species, C, across all *M* subjects that have been categorized in at least one instance for the two sets ℛ^(*ν,T* −1)^ and 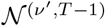, pertaining to different subjects. The cross-subject re-establishment empirical probability is then expressed 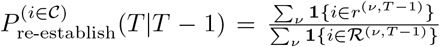, where **1**{condition} denotes the indicator function. Similarly, the probability 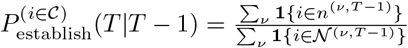 is defined.

### Permutation test: cross-subject re/establishment probability analysis

Given paired values 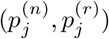, where 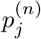, and 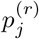 are the per-case empirical probability estimate for establishment, and re-establishment of a given species respectively. We computed the differences 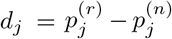 and excluded pairs with *d*_*j*_ = 0. The observed test statistic was defined as the sum of differences, *T* ^(obs)^ = ∑_*j*_ *d*_*j*_. To construct the null distribution, we fixed the magnitudes |*d*_*j*_| and randomly assigned an independent sign ±1 to each difference. Specifically, for each permutation *b* = 1, …, *B*, we sampled signs 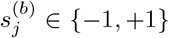 with equal probability and formed 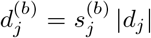, and 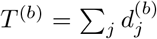. A one-sided *p*-value was computed as 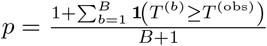, where **1**{condition} is the indicator function.

### Similarity Correlation

To conduct the ‘similarity correlation’ analysis for a specific test subject *ν*, we followed these steps: We first define the set *q* = *u* ∪ *s* ∪ *r*^(*t*≤*τ*)^. Additionally, we define the set of ‘new species’ present at the recovery endpoint as 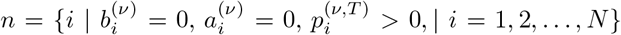. Next, we define **c** := (*c*_1_, …, *c*_*M*_)^T^, where 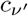 represents the species richness of the reference baseline sample 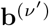. We then calculate *M* similarity values between the vector 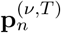 and the *M* reference baseline samples in the subspace *n*, 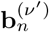, using Jaccard similarity. We define the vector containing these similarity measures as 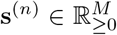. Similarly, we compute 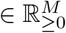 using the set *q*. The similarity Correlation, *sCorr*, is expressed as:

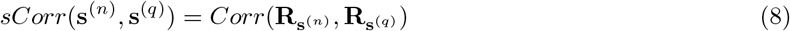

Where

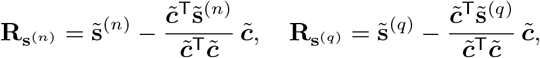

and the tilde denotes mean-centering across the *M* reference baselines (e.g., 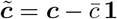, with 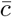 the mean of the elements of ***c***). Thus, 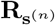 and 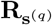 are the residuals of ordinary least-squares regressions of **s**^(*n*)^ and **s**^(*q*)^ on ***c*** with an intercept, and *Corr*(·, ·) denotes the Pearson correlation.

Equivalently, *sCorr* is the partial Pearson correlation between **s**^(*n*)^ and **s**^(*q*)^ controlling for the reference-baseline richness vector **c**, obtained by linearly regressing each of **s**^(*n*)^ and **s**^(*q*)^ on **c** (with an intercept) and correlating the residuals (`pcor.test` from the `ppcor` R package). Its significance was assessed with the two-sided *t*-test for a partial correlation, 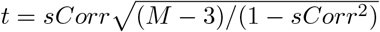, referred to a *t* distribution with *M* − 3 degrees of freedom (*M* reference baselines, minus two, minus one conditioning variable). The persubject p-values were adjusted across the subjects of each dataset with the Benjamini–Hochberg procedure at a false discovery rate of *α* = 0.05 (Fig. 5**e**); a subject was counted as showing a positive, significant *sCorr* when the adjusted p-value was below *α* and *sCorr >* 0.

### Population dynamics model

To simulate the effect of interspecies interactions on colonization via historical contingency mechanism, we used the generalized Lotka–Volterra (GLV) model, which describes the dynamics of *N* interacting species as a system of ordinary differential equations,

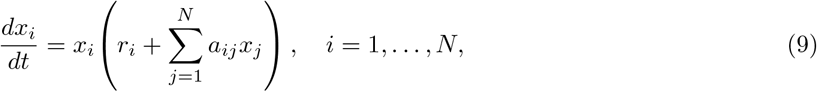

where *x*_*i*_ denotes the population density of species *i, r*_*i*_ is the intrinsic growth rate of species *i, a*_*ij*_ represents the effect of species *j* on species *i*, and *a*_*ii*_ corresponds to self-interaction (logistic) terms, with *a*_*ii*_ *<* 0 for all *i*.

### Historical contingency simulation

To illustrate the historical contingency mechanism using the GLV model, we conducted the following simulation: We generated the off-diagonal elements of the matrix 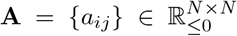 from −|*Z*|, where *Z* ∼ *N* (*µ, σ*^2^), with a probability of *c*, and set them to 0 with a probability of 1 − *c*. The diagonal elements of **A** were set to −*d*, where *d >* 0. We defined the intrinsic growth rate vector 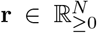 with *r*_*i*_ = 1 for all *i*. Next, we generated a matrix 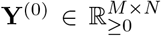, where each row contained *k* non-zero values chosen randomly and independently from *U* [0, *K*], with *K >* 0. We then determined the steady state matrix 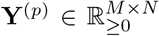, normalizing **Y**^(*p*)^ such that: 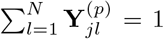 for all *j*. Subsequently, we set values smaller than *η* (the ‘minimum abundance threshold’) to 0 and renormalized **Y**^(*p*)^. We defined **y** as one of the rows in **Y**^(*p*)^, and for all rows in **Y**^(*p*)^, we replaced zeros with a small *ϵ* (an immigration rate, a minor influx from the regional pool), and found the new steady state 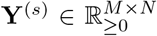, normalizing the rows. The new steady state of **y** is denoted as **y**^(*s*)^. Afterward, we set values smaller than *η* to 0 and renormalized **Y**^(*s*)^. We defined 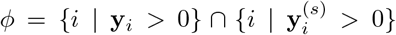 and 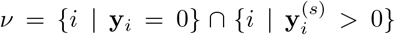. Finally, we applied the similarity correlation analysis, calculating the *sCorr* value using the sample **y**^(*s*)^ as the test sample and the other samples in **Y**^(*s*)^ as the reference. In our simulations, we used the parameters: (*N, M, µ, σ, c, d, k, K, η, ϵ*) = (200, 300, 0, 5, 0.05, 1, 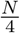, 1, 10^−4^, 10^−4^). For numerical integration, we employed the `solve ivp` function from the `scipy.integrate` library in Python, utilizing the Runge-Kutta method of order 5(4). The condition for a steady state was defined as max_*i*_ 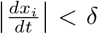, where *δ* = 10^−5^. If the steady state condition was not satisfied by *t* = 1000, the sample was rejected.

To demonstrate that the interactions among species are responsible for the historical contingency effect, we employ the aforementioned procedure. Prior to determining **y**^(*s*)^, we define *ξ* = {*i* | **y**_*i*_ *>* 0} and establish the following:

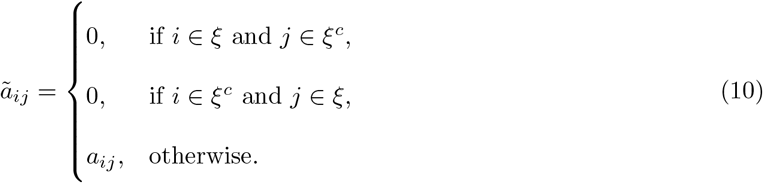

Subsequently, we utilize **Ã** to obtain **y**^(*s*)^.

### Similarity functions

Consider two microbial samples, each represented by the species abundance vectors **u** = (*u*_1_, *u*_2_, …, *u*_*N*_) and **v** = (*v*_1_, *v*_2_, …, *v*_*N*_), where *u*_*i*_ and *v*_*i*_ denote the relative abundances of species *i*, and *N* is the total number of species considered. For convenience, define the index sets of detected species as *U* = {*i* ∈ {1, …, *N* } | *u*_*i*_ *>* 0} and, *V* = {*i* ∈ {1, …, *N* } | *v*_*i*_ *>* 0}. We define the following similarity measures:

1. Root Jensen-Shannon divergence similarity (rJSD):

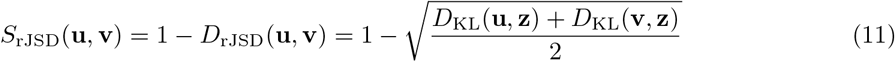

where **z** ≡ (**u** + **v**)*/*2, and 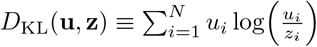 is the Kullback–Leibler divergence between **u** and **z**.
2. Jaccard similarity:

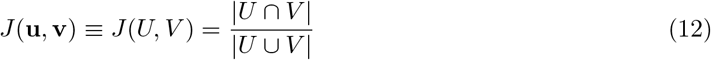
3. Specificity similarity:

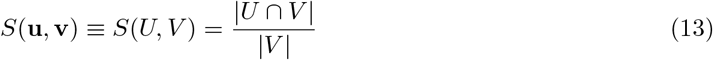

Here, | · | denotes the number of elements of a set.

### Lost and newly detected species

The ‘lost species’ are baseline taxa that were not detected again at any later point, neither during treatment nor at any follow-up sample,

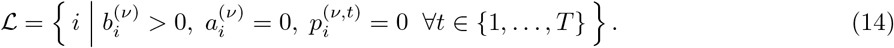

The remaining two categories concern taxa that were never observed at baseline. Let

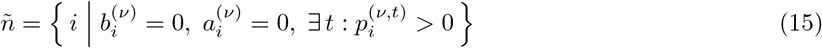

denote all taxa first detected during the follow-up period. Within *ñ*, the ‘colonizers’ are those established at the end of the study, in the sense of being present in the final two follow-up samples,

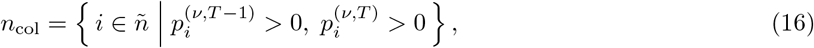

while the ‘transient’ taxa are those detected in exactly one follow-up sample and absent from the final one,

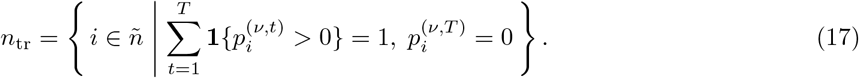

The two sets are disjoint but do not exhaust *ñ*; taxa detected more than once yet absent from the final sample belong to neither, and are excluded from the comparison between them.

### Functional Profiles

Consider a local community comprising *N* species denoted as [*x*_1_, …, *x*_*N*_], where *x*_*i*_ represents the relative abundance of species *i*. Additionally, consider *P* functions represented as [*f*_1_, …, *f*_*P*_], where *f*_*α*_ denotes the relative abundance of function *α*. Each species *x*_*i*_ is associated with a corresponding vector of functional copy numbers 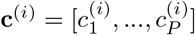, such that 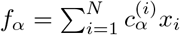.

### Functional Similarity Measures

To determine the functional similarity *s*_*ij*_ = *J*_*W*_ (**c**^(*i*)^, **c**^(*j*)^) between taxon-*i* and taxon-*j*, we employ the weighted Jaccard similarity metric applied to the genomes of the two taxa:

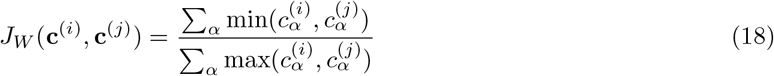

### AUC analysis: discrimination between colonizing and transient new species

For each subject we compared two groups of scores: the mean ‘functional similarity’ to the reference set of the colonizers, denoted 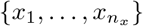, and the corresponding values for the transient taxa, 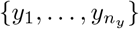 (defined above). Non-finite values were removed before testing. The separation between the two groups was quantified by the area under the receiver operating characteristic curve (AUC), computed with `roc auc score` from the `sklearn.metrics` module, using the raw scores as the discriminant variable and labelling the colonizers as the positive class and the transient taxa as the negative class. No threshold on the score is selected: the AUC integrates over all possible cut-offs and is equal to

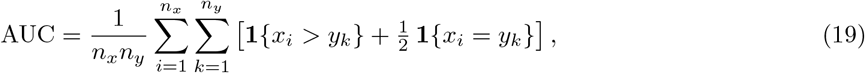

the probability that a randomly chosen colonizer has a higher mean functional similarity than a randomly chosen transient species, with tied pairs contributing one half. An AUC of 0.5 indicates no discrimination, and values above 0.5 indicate that the colonizers are the more functionally similar group.

Statistical significance was assessed with the two-sided Mann–Whitney *U* test, using `mannwhitneyu` from the `scipy.stats` module with `alternative=“two-sided”` and `method=“auto”`, which selects the exact null distribution for small samples without ties and the tie-corrected normal approximation otherwise. The two quantities describe the same comparison, since AUC = *U/*(*n*_*x*_*n*_*y*_) where *U* is the Mann-Whitney statistic for the colonizers; the AUC is therefore reported as the effect size and the Mann-Whitney test as its significance. Raw p-values were corrected for multiple comparisons across subjects using the Benjamini-Hochberg procedure at a false discovery rate of *α* = 0.05. Subjects for which either group contained fewer than *n*_min_ species were not tested and were excluded from the correction family, so that the procedure was applied only to the subjects actually tested.

### Data and pre-processing

The microbiome data analyzed in this study are sourced from previously published research. Throughout the presence-absence analyses, a taxon was considered present when its abundance was greater than zero (after rarefaction, where applied) and absent when no reads were detected. Thus, “absence” refers to non-detection at the applied sequencing depth rather than definitive biological absence. (1) Palleja et al. [6] utilized microbiome taxonomic data represented as relative abundance profiles of species-level taxa (mOTUs) obtained through shotgun metagenomic sequencing. The raw reads underwent standard quality-control procedures and were mapped to a reference database of universal marker genes to generate mOTU-based taxonomic profiles. We rarefied the data to account for variation in sequencing depth. Subjects who were not sampled at all study time points were excluded from the longitudinal analysis; however, their baseline samples were retained as reference samples where relevant. (2) Sewunet et al. [5] employed microbiome taxonomic data derived from 16S rRNA gene sequencing, represented as compositional abundance profiles of operational taxonomic units (OTUs). The raw reads were processed and clustered using VSEARCH against the Greengenes database (97% similarity threshold) to obtain OTUs and their taxonomic assignments. We rarefied the data to account for differences in sequencing depth. One subject was excluded from the longitudinal analysis because its species richness during antibiotic treatment exceeded its baseline by more than twofold, whereas all other subjects had higher richness at baseline than during treatment. Its baseline sample was retained as a reference because its richness was comparable to that of the other baseline samples. (3) Messaoudene et al. [32] conducted a study utilizing microbiome taxonomic data obtained from 16S rRNA gene sequencing. The paired-end 16S rRNA sequencing reads were processed using QIIME 2 [43] (amplicon distribution, version 2025.4). Raw FASTQ files were imported via a manifest file and underwent quality control and denoising through DADA2 [44], resulting in the generation of an ASV feature table and representative sequences. Taxonomic classification was performed using the QIIME 2 feature-classifier [45] ‘classify-sklearn’ method with a pre-trained Naive Bayes classifier (scikit-learn 1.4.2; 2024.09 backbone reference database). Functional profiles were inferred from the 16S rRNA sequencing data using PICRUSt2 [46]. The taxonomic data were rarefied. Subjects in the antibiotic-only groups who were not sampled at all study time points were excluded from the longitudinal analysis, although their baseline samples were retained as references where relevant. Baseline samples from participants in the other treatment groups were also included as reference healthy configurations. (4) For Yaffe et al. [13], we processed the raw paired-end shotgun metagenomic reads ourselves. Reads were quality-checked with FastQC and MultiQC [47], then quality-filtered and depleted of host-derived sequences (human hg38) with KneadData, and only retained read pairs were used downstream. Species-level relative-abundance profiles were generated with MetaPhlAn 4 [48] (database `mpa vOct22 CHOCOPhlAnSGB 202403`) and merged across samples, retaining species-level entries only. These profiles were not rarefied. Functional profiles (gene families, reactions, and pathways) were generated with HUMAnN 4 [49] (ChocoPhlAn v4 alpha SGB pangenome database and EC-filtered UniRef90 protein database, version 2019 06) from the filtered reads and the MetaPhlAn profiles, and were kept in the original HUMAnN output units.

## Code availability

Codes used in this work are available at https://github.com/ShayaKahn/Recovery_from_antibiotics.

