## Supplementary Information for "Individual-specific recovery of the healthy human gut microbiota after antibiotics"

#### Contents


**1 Evaluation of the criteria for individual-specific recovery** **2**

**2 Antibiotic-associated disruption and the ciprofloxacin-treated cohort** **3**

**3 Individual-specific recovery without standardization** **4**

**4 Individual-specific recovery based on phylogenetic similarity** **5**

**5 Taxonomic resolution in surrogate-data analysis of post-antibiotic recovery** **6**

**6 The relation between species assemblage and abundance** **9**

**7 The effect of metadata on individual-specific recovery** **10**

**8 Rate of accumulation of persistent newly detected taxa** **11**

**9 Correlation between surviving probability and average return time** **14**

**10 Antibiotic-associated depletion and directed switching toward a specific alternative state** **15**

**11 Cross-species/subject re/establishment analysis for different abundances** **16**

**12 Historical contingency simulation using a consumer-resource model** **18**

**13 Functional association between new colonizers and baseline species in shotgun metagenomic data** **20**

### 1 Evaluation of the criteria for individual-specific recovery

In the main analysis, a subject is classified as exhibiting individual-specific recovery when two criteria are satisfied: (i) the subject's own baseline ranks first among all reference baselines (rank = 1), and (ii) the own-baseline similarity is significantly greater than expected from the surrogate distribution. The rank = 1 criterion becomes increasingly stringent as the number of available reference baselines increases because the recovered community must be more similar to its own baseline than to every alternative baseline. We therefore additionally evaluated individual-specific recovery using only the significance criterion, which is less directly dependent on the number of available reference baselines. Supplementary Fig. 1 compares the fraction of subjects satisfying both criteria with the fraction satisfying the significance criterion alone across different reference-cohort sizes, providing a complementary assessment of the robustness of individual-specific recovery to the number of available reference baselines.

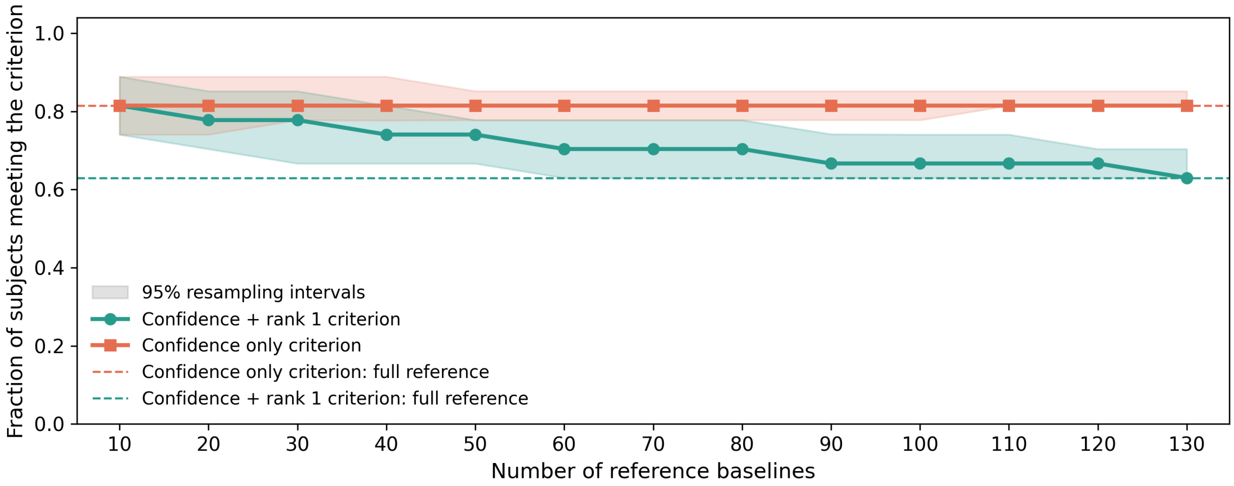


**Supplementary Figure 1: Sensitivity of the criteria for individual-specific recovery to reference-cohort size.** For each focal subject, eligible reference baselines, excluding that subject, were randomly sampled without replacement at cohort sizes ranging from 10 to 130. Within each of 100 repetitions, reference subsets were nested across cohort sizes by taking progressively larger portions of the same random permutation. Surrogate data analysis was performed using rJSD similarity with all taxa included. Lines show the median fraction of subjects satisfying both the rank = 1 and Benjamini–Hochberg-adjusted $p < 0.05$ criteria (green), or the adjusted $p$-value criterion alone (orange). Shaded regions represent 95% resampling intervals, and dashed lines show the corresponding results obtained using all eligible reference baselines.

### 2 Antibiotic-associated disruption and the ciprofloxacin-treated cohort

#### Measured disruption across cohorts

The main text groups the four cohorts by the impact expected from their antibiotic protocols. Here we report two measures of the disruption actually observed. The mean Bray-Curtis dissimilarity between the final antibiotic sample and the subjects' own baselines was 0.647 for Yaffe et al. [1], 0.810 for Sewunet et al. [2], 0.839 for Messaoudene et al. [3], and 0.957 for Palleja et al. [4], reproducing the ordering anticipated from the protocols. The mean retention of baseline species was 47.1%, 19.4%, 25.6%, and 34.8% for the same four cohorts, respectively. Retention likewise identifies the Yaffe et al. cohort as the least disrupted, but does not reproduce the ordering among the remaining three: the Palleja et al. cohort retained a larger fraction of baseline taxa (34.8%) than the Messaoudene et al. (25.6%) and Sewunet et al. (19.4%) cohorts, despite showing the largest compositional shift. The two measures need not coincide, because Bray-Curtis dissimilarity is abundance-weighted whereas retention is a detection-based count that is additionally sensitive to sequencing depth and feature resolution, both of which differ across the four studies. We therefore use Bray-Curtis dissimilarity to compare measured disruption between cohorts, and the per-subject fraction of undetected baseline taxa (Fig. 2**h** of the main text) for comparisons within them.

#### Individual-specific recovery in the ciprofloxacin-treated cohort

Supplementary Fig. 2 presents the surrogate-data analyses using rJSD and Jaccard similarity, together with the null-model analysis, for the ciprofloxacin-treated cohort of Yaffe et al. [1]. These panels are the counterparts of Fig. 2**b**, **d**, and **g** of the main text and are reported here because this cohort is substantially larger than the others, so its strip plots do not fit alongside them in the main figure.

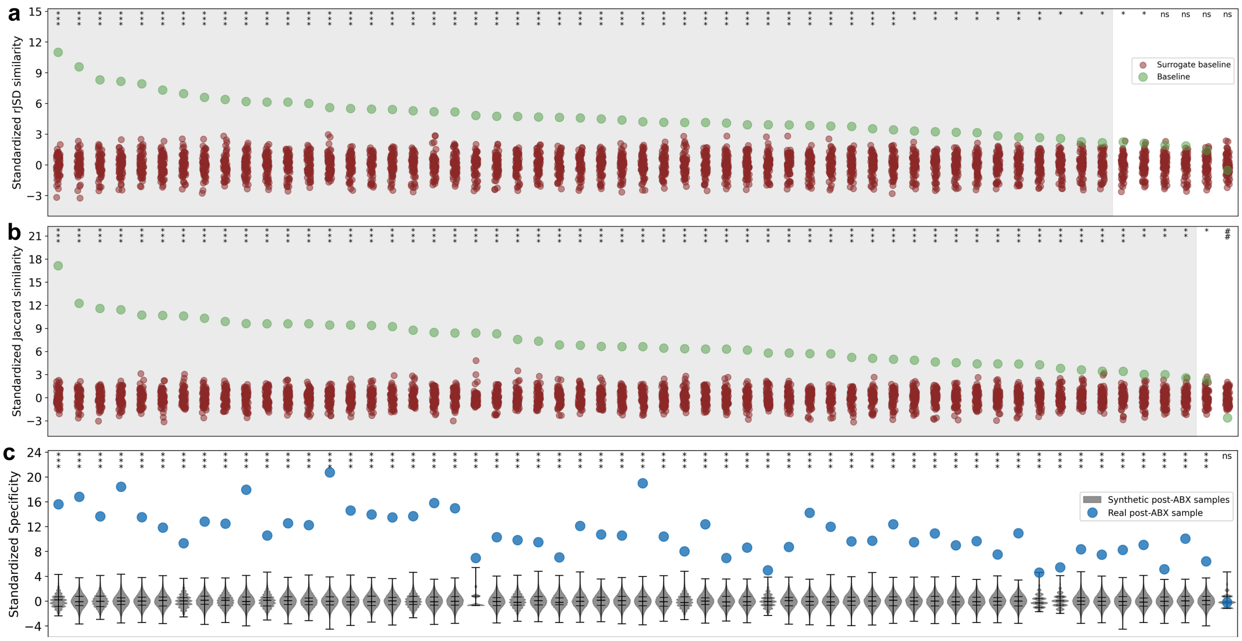


**Supplementary Figure 2: Individual-specific recovery in the ciprofloxacin-treated cohort. a-c**, Analyses parallel to Fig. 2**b**, **d**, and **g** of the main text, for the antibiotic-treated subjects of Ref. [1] ($n = 57$). **a**, Surrogate-data analysis using rJSD similarity with all taxa included: 93% (53 of 57) of subjects met the significance criterion, and 89% (51 of 57) additionally met the dual criterion. **b**, The same analysis using Jaccard similarity with the survived species excluded: 96% (55 of 57) of subjects met the dual criterion. **c**, Null-model analysis: 98% (56 of 57) of subjects showed significantly greater specificity than expected under the null model. Symbols and significance notation follow Fig. 2 of the main text.

### 3 Individual-specific recovery without standardization

In the main text, Fig. 2**b** presents individual-specific recovery using rJSD similarities standardized relative to the corresponding surrogate distributions. Here, we repeat the same analysis using the raw rJSD similarity values, without standardization, to show the absolute similarity between each post-antibiotic community and its own baseline relative to the alternative baseline communities (Supplementary Fig. 3).

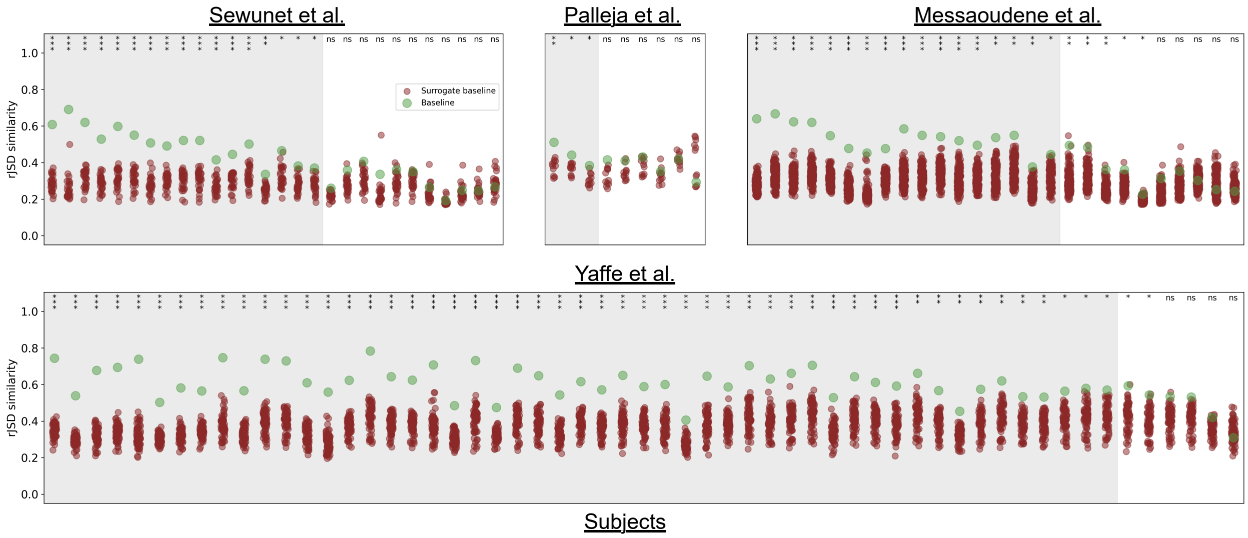


**Supplementary Figure 3: Individual-specific recovery based on unstandardized rJSD similarity.** The analysis presented in Fig. 2**b** of the main text is shown using the raw rJSD similarity values, without standardization relative to the corresponding surrogate distributions.

### 4 Individual-specific recovery based on phylogenetic similarity

To determine whether individual-specific recovery is also observed when phylogenetic relationships among taxa are considered, we repeated the surrogate-data analysis using unweighted UniFrac similarity, calculated as one minus the unweighted UniFrac distance. The analysis was performed for subjects in the Messaoudene et al. cohort (Ref. [3]) using both similarities standardized relative to the corresponding surrogate distributions and the unstandardized raw values (Supplementary Fig. 4**a**,**b**, respectively). The results were consistent with those obtained using rJSD similarity: 74% of subjects satisfied both the rank = 1 and significance criteria. Thus, individual-specific recovery remains evident when similarity is defined based on phylogenetic community composition.

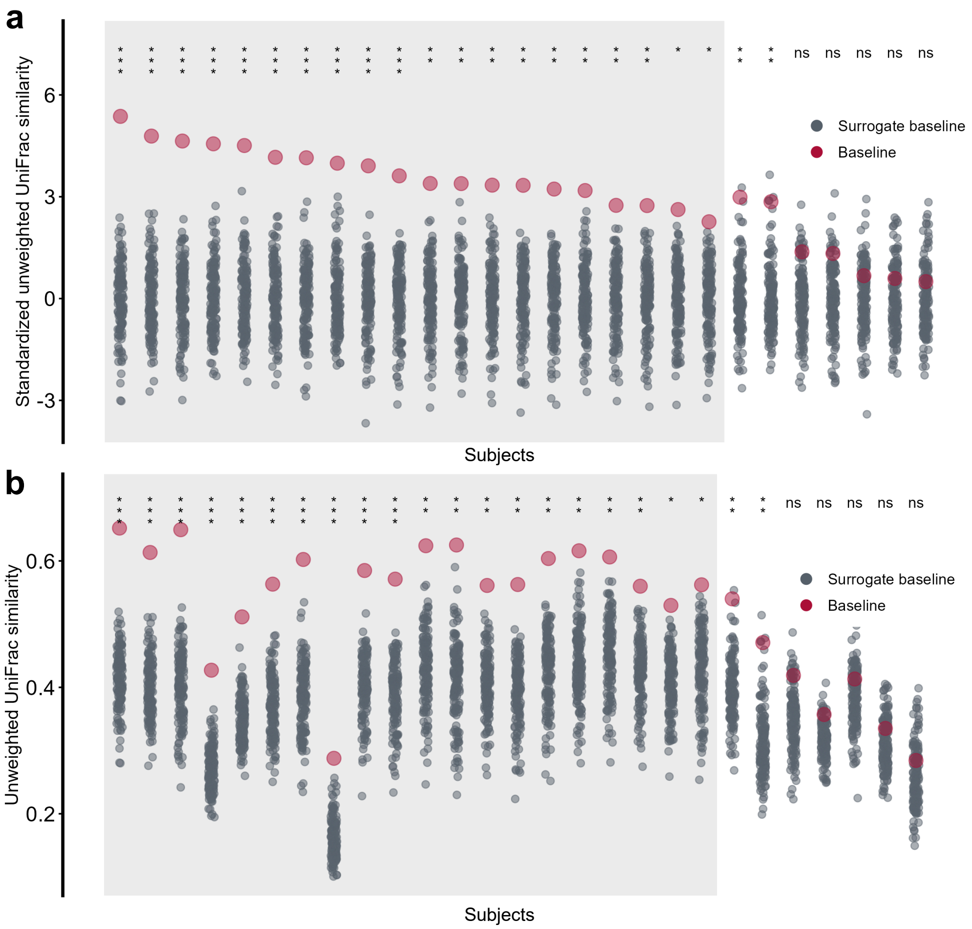


**Supplementary Figure 4: Individual-specific recovery based on unweighted UniFrac similarity.** **a**, Surrogate-data analysis analogous to that presented in Fig. 2**b** of the main text, performed for subjects in the Messaoudene et al. cohort (Ref. [3]) using unweighted UniFrac similarity, calculated as one minus the unweighted UniFrac distance. Similarities are standardized relative to the corresponding surrogate distributions. **b**, The same analysis shown using the unstandardized raw unweighted UniFrac similarity values.

### 5 Taxonomic resolution in surrogate-data analysis of post-antibiotic recovery

The individual-specific recovery pattern observed at fine taxonomic resolution disappeared when communities were analyzed at the genus level (Supplementary Fig. 5a,b). In both datasets, genus-level surrogate data analysis no longer showed the preferential similarity of post-antibiotic communities to their own baselines seen at the ASV or OTU level. The trajectories and taxon-overlap comparisons for subjects 18 AMC and 15 AMC illustrate how genus-level aggregation can obscure differences in community membership and dynamics

(Supplementary Fig. 5c,d). These results suggest that individual-specific recovery is reflected primarily in the composition of taxa within genera, highlighting the importance of taxonomic resolution when assessing host-specific community recovery.

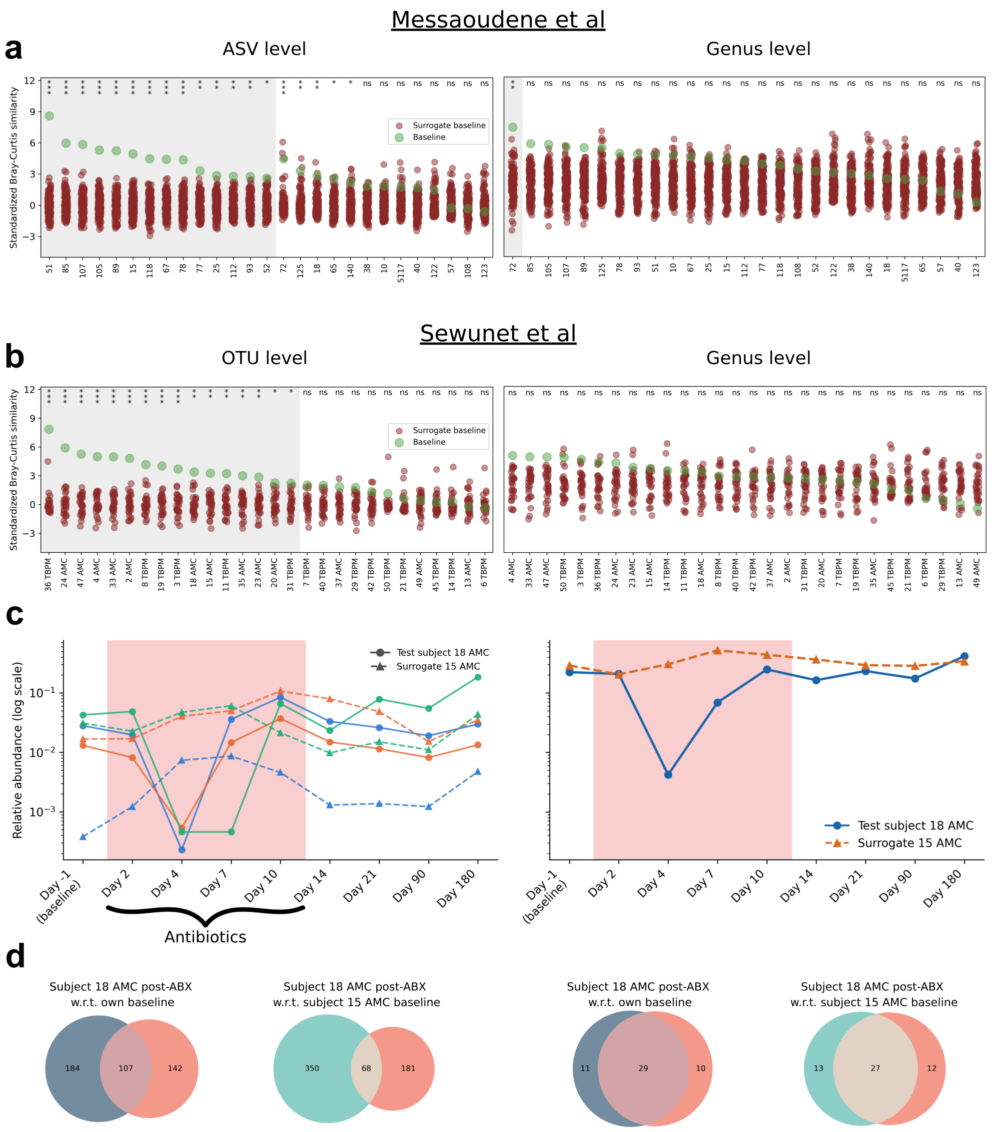


**Supplementary Figure 5: Surrogate-data analysis and community recovery at different taxonomic resolutions. a-b**, Surrogate-data analysis using Bray-Curtis similarity for the Messaoudene et al. dataset (**a**; ASV level, left; genus level, right) and the Sewunet et al. dataset (**b**; OTU level, left; genus level, right). For each subject, the final post-antibiotic community was compared with its own baseline (green) and the baselines of other subjects (red). **c**, Relative-abundance trajectories for test subject 18 AMC and surrogate subject 15 AMC in the Sewunet et al. dataset. The left plot shows individual OTUs within the selected genus that were detected at every sampled timepoint in both subjects; colors identify OTUs. The right plot shows the total relative abundance of the genus, obtained by summing all its mapped OTUs, including those not continuously detected. Solid lines with circles represent the test subject, and dashed lines with triangles represent the surrogate. Abundances are displayed on a logarithmic scale; sampling days are equally spaced for visualization, and pink shading marks antibiotic treatment. **d**, Taxon overlap between the final post-antibiotic community of subject 18 AMC and either its own baseline or the baseline of subject 15 AMC, at the OTU level (left) and genus level (right). Dark blue and turquoise circles represent the test and surrogate baselines, respectively; orange circles represent the test subject's final post-antibiotic community. Numbers indicate taxon counts in the exclusive and shared regions.

### 6 The relation between species assemblage and abundance

Our analysis, as depicted in Fig. 2**d** of the main text, utilizes the Jaccard similarity, a measure based on presence-absence data. Given that the analysis involves comparing samples after selecting specific subsets of species, the direct application of an abundance-based similarity measure may present challenges [5]. To investigate whether assemblage similarity correlates with abundance similarity, we have defined two complementary metrics. These metrics are based on those introduced in Ref. [6].

First, 'unweighted similarity' measures the similarity between two microbial samples based only on species presence-absence:

$$\textit{Unweighted similarity} = J(\mathbf{x}, \mathbf{y}), \tag{1}$$

where $J(\cdot, \cdot)$ denotes the Jaccard similarity, and $\mathbf{x}, \mathbf{y} \in \mathbb{R}^N_{\geq 0}$ represent two microbial samples with $N$ species.

Second, 'weighted similarity' measures the abundance-based similarity between the species shared by the two samples, after renormalizing their abundances within the shared set:

$$\textit{Weighted similarity} = 1 - D_{rJSD}(\hat{\mathbf{x}}, \hat{\mathbf{y}}), \tag{2}$$

where

$$\hat{\mathbf{x}} = \frac{(x_i)_{i \in S}}{\sum_{j \in S} x_j}, \qquad \hat{\mathbf{y}} = \frac{(y_i)_{i \in S}}{\sum_{j \in S} y_j},$$

and

$$S = \big\{ i \in \{1, \ldots, N\} \mid x_i > 0,\ y_i > 0 \big\}.$$

In Supplementary Fig. 6, we observed a positive correlation between unweighted similarity and weighted similarity when comparing each subject's baseline sample to their post-antibiotic sample. This suggests that greater recovery of species composition is typically accompanied by greater recovery of the relative abundances of shared species.

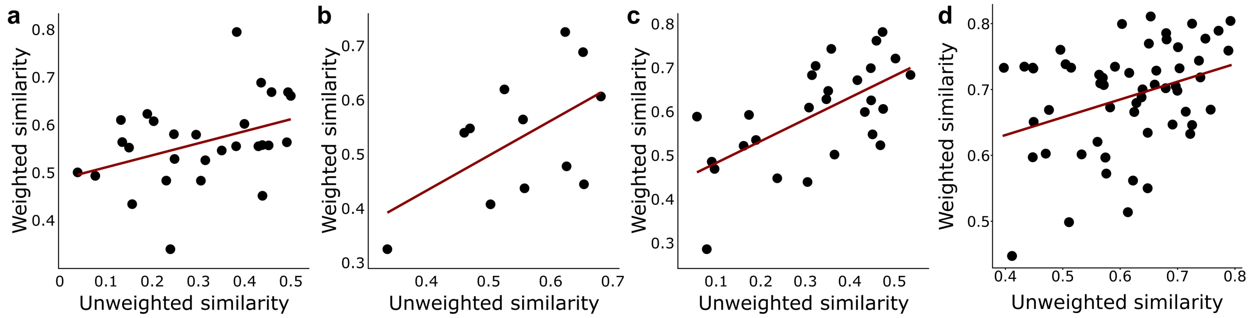


**Supplementary Figure 6: a-d**, Correlation between the unweighted similarity (Jaccard similarity) and the weighted similarity (one minus rooted Janson-Shannon divergence calculated between the renormalized abundances of the shared species) measured between in the pre- and post-antibiotics samples, for subjects from Ref. [2] (**a**, $n = 28$, $r = 0.39$, $p = 0.041$), Ref. [4] (**b** $n = 9$, $r = 0.549$, $p = 0.065$), Ref. [3] (**c** $n = 27$, $r = 0.623$, $p = 5 \cdot 10^{-4}$), and Ref. [1] (**d** $n = 57$, $r = 0.35$, $p = 0.008$).

### 7 The effect of metadata on individual-specific recovery

In the main text, we report the fraction of subjects exhibiting individual-specific recovery in each cohort, according to the abundance-based surrogate data analysis and its complementary presence-absence version, and show using a binomial test that these fractions are unlikely to arise by chance under the corresponding null expectation. An alternative explanation, however, is that microbiomes recover not toward truly individual-specific states, but toward states characteristic of broader subgroups of individuals. Such subgroups could reflect shared factors such as sex, age, BMI, geographic location, ethnicity, diet, or other host characteristics. Recovery toward subgroup-specific states could therefore generate an apparent preference for the original baseline relative to microbiomes sampled from the cohort as a whole.

To examine this possibility, Supplementary Fig. 7 shows a PCoA based on $rJSD$ distances between baseline microbiomes from Ref. [3], colored according to sex, age, and BMI. PERMANOVA revealed no significant association between baseline microbiome composition and any of these factors. Moreover, the available metadata indicate that this cohort is relatively homogeneous; for example, all participants were recruited from the same city and country. These results provide no evidence that the observed individual-specific recovery is driven by broad subgroups defined by the measured host characteristics, although effects of unmeasured or more subtle covariates cannot be excluded.

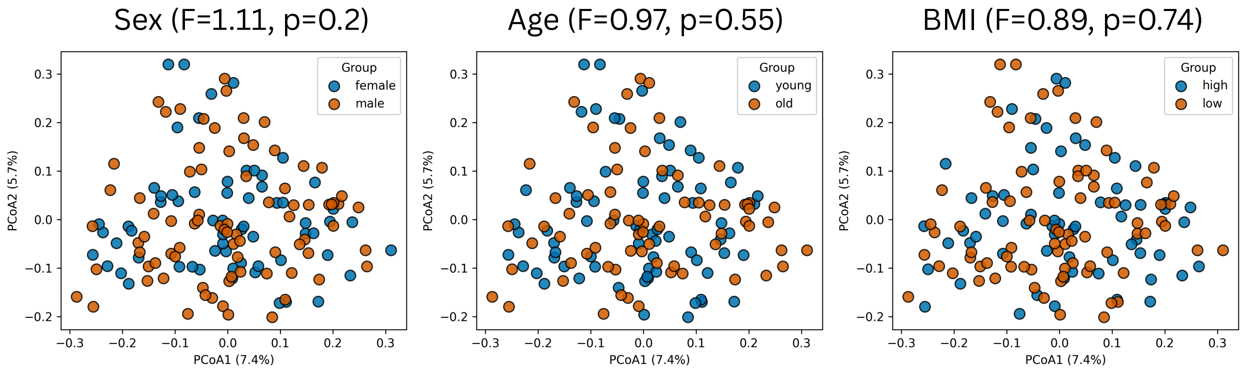


**Supplementary Figure 7:** PCoA visualization of subjects from Ref. [3] based on rJSD distances. Subjects are colored according to three metadata variables: sex, age group, and BMI group. Left: subjects are divided by sex; PERMANOVA shows no significant separation between groups ($F = 1.11$, $p = 0.20$). Middle: subjects are divided into young and old groups, defined as below and above the cohort mean age, respectively; PERMANOVA shows no significant separation ($F = 0.97$, $p = 0.55$). Right: subjects are divided into low-BMI and high-BMI groups, defined as BMI $<$ 25 and BMI $>$ 25, respectively; PERMANOVA shows no significant separation ($F = 0.89$, $p = 0.74$). PERMANOVA tests were performed using the `permanova` function from `skbio.stats.distance`.

### 8 Rate of accumulation of persistent newly detected taxa

In the main text, we show that new taxa accumulate gradually over weeks to months and that their mean arrival time depends on the duration of the observation window. To further characterize this process, we quantified the rate at which persistent newly detected taxa emerged following antibiotic treatment. Here, a newly detected taxon was considered persistent if, after its first detection following treatment, it remained detectable at every subsequent follow-up sample; taxa detected only briefly or intermittently were considered non-persistent. This persistence criterion is specific to the present analysis and is not the colonizer/transient classification used in the functional analysis (Methods), which is anchored on the final two follow-up samples.

Supplementary Fig. 8 shows two main patterns. First, the first-detection rate of persistent new taxa was higher in antibiotic-treated subjects than in the corresponding control subjects. Second, among antibiotic-treated subjects, the first-detection rate generally decreased over time but remained above zero at the latest follow-up for which persistence could be evaluated. These results are consistent with a prolonged post-antibiotic assembly process that gradually slows over time rather than exhibiting a sharply defined completion point. Consequently, summary measures such as mean arrival time are inherently influenced by study duration and sampling schedule, motivating their comparison primarily within studies rather than across cohorts with different observation windows.

Persistence among newly detected taxa was uncommon. In the Messaoudene et al. dataset [3], participants had a mean of $158.4 \pm 46.6$ newly detected taxa before the final follow-up, of which only $19.0 \pm 10.4$ were persistent by the criterion above. In the Sewunet et al. dataset [2], $174.1 \pm 59.6$ taxa were newly detected and only $33.8 \pm 20.6$ were persistent. Most newly detected taxa therefore did not persist to the end of follow-up, indicating that establishment in the recovering gut community is a selective process. Values represent the mean $\pm$ sample standard deviation across participants; taxa first detected at the final follow-up were excluded.

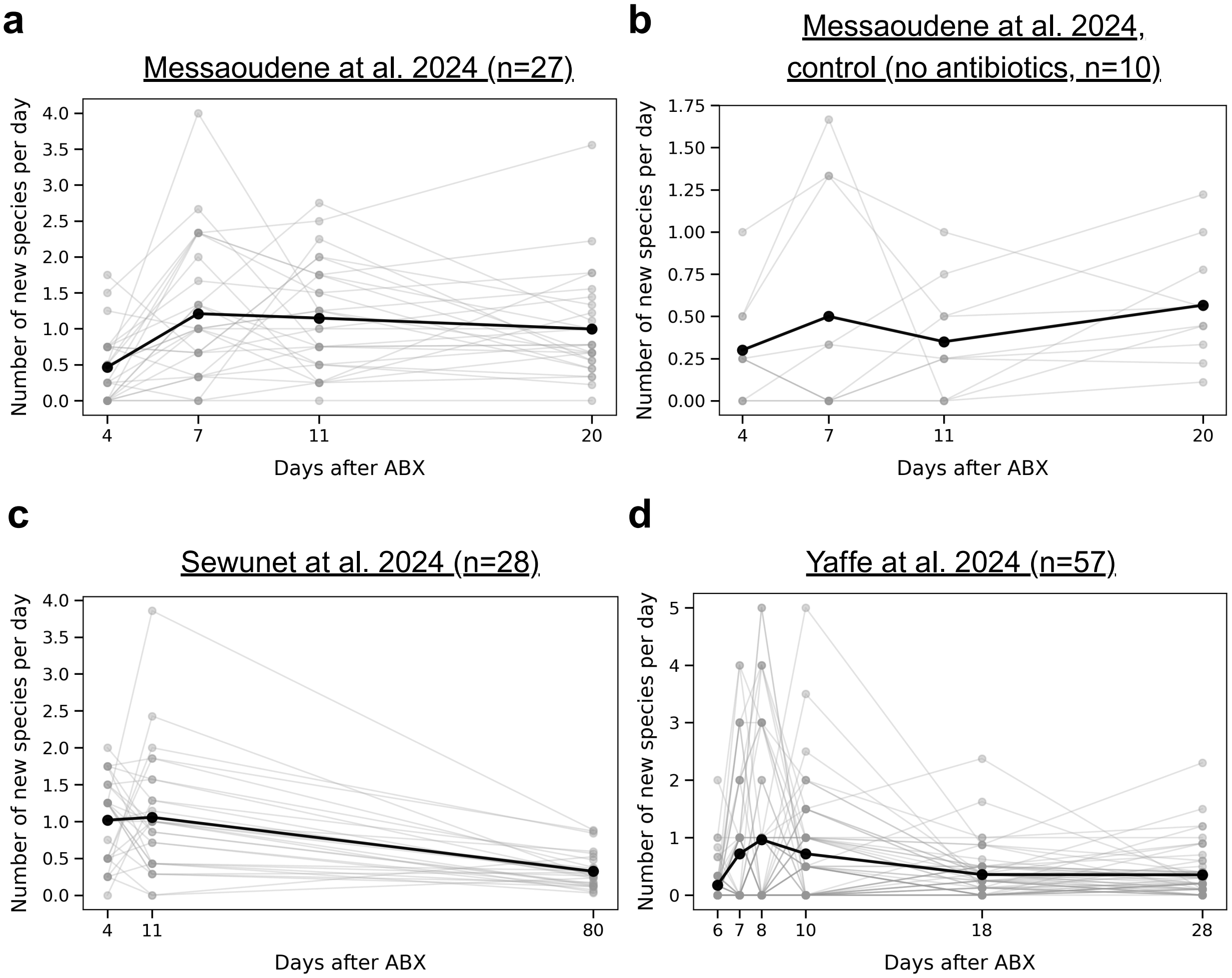


**Supplementary Figure 8: Rate of accumulation of persistent newly detected taxa following antibiotic treatment**. The number of persistent new species appearing per day is shown for **a)** the antibiotic-treated cohort from Messaoudene et al. [3], **b)** the corresponding control cohort, **c)** the antibiotic-treated cohort from Sewunet et al. [2], and **d)** the antibiotic-treated cohort from Yaffe et al. [1]. A species was classified as new at a given time point if it was undetected at baseline and during antibiotic treatment, or at the corresponding sampling period in controls, was first detected at that time point, and remained detected at all subsequent follow-ups. Counts were divided by the number of days since the preceding sample to obtain an emergence rate. Grey points and connecting lines represent individual participants, whereas black points and lines show the cohort mean. The final follow-up was excluded because persistence after first detection could not be evaluated.

### 9 Correlation between surviving probability and average return time

To enhance the association between early return time and the classification of early-returning species as survived species, we introduce two distinct metrics: (1) 'probability to survive' and (2) 'average return time'. The probability to survive for a particular species is defined as follows:

$$p_S = \frac{\text{\# of subjects where this species classified as survived}}{\text{\# of the total classification cases}}. \tag{3}$$

Similarly, the probability of returning on day $D^{(t)}$ is expressed as:

$$p_t = \frac{\text{\# of subjects where this species classified as returned at day } D^{(t)}}{\text{\# of the total classification cases}}. \tag{4}$$

The average return time is defined as follows:

$$\bar{t} = \sum_t D^{(t)} \cdot P(\text{Returned at day } D^{(t)} | \text{Returned}) \tag{5}$$

Supplementary Fig 9 illustrates a significant correlation across three distinct datasets between the probability to survive and the average return time. This finding suggests that species with a higher likelihood of survival across subjects tend to return earlier across different subjects.

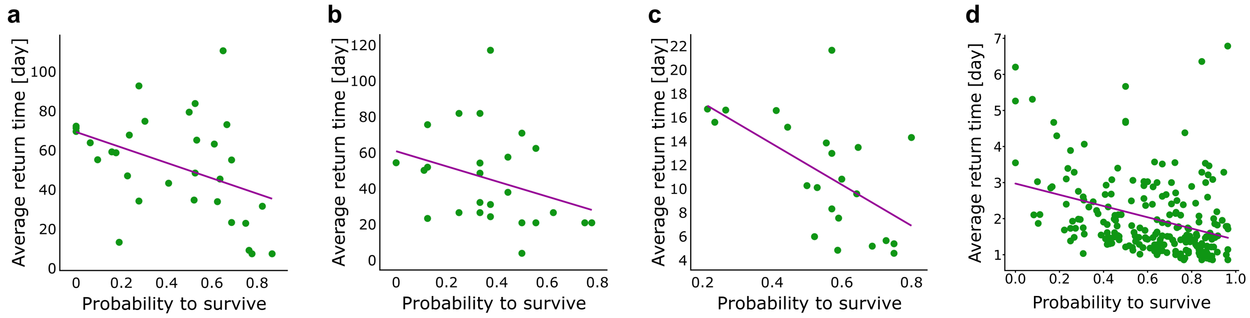


**Supplementary Figure 9: a–d**, For each subject, each species was classified as having survived antibiotic administration, returned at day $D^{(t)}$ after antibiotic administration, or neither. The panels show negative Spearman correlations between species' probability to survive and the average return time: Ref. [2] (**a**; $\rho = -0.46$, $p = 0.009$), Ref. [4] (**b**; $\rho = -0.41$, $p = 0.03$), Ref. [3] (**c**; $\rho = -0.62$, $p = 0.002$), and Ref. [1] (**d**; $\rho = -0.37$, $p = 2.04 \times 10^{-8}$). Analyses include species with classification frequencies exceeding 14, 7, 13, and 10, respectively. Across all panels, species whose average return time is based on a single case are excluded.

### 10 Antibiotic-associated depletion and directed switching toward a specific alternative state

In each of the two analyzed datasets [2,3], exactly four subjects showed standardized own-baseline specificity significantly below the random-sampling null distribution (Fig. 4**b** in the main text). The null model represents arbitrary restoration from the cohort-wide baseline pool; these subjects were therefore less similar to their own baselines than expected even under arbitrary restoration. This lower-tail result indicates a directed, non-random departure from the original state, consistent with a transition toward a specific alternative configuration rather than random drift. These cases were uncommon and do not imply that every subject lacking significant individual-specific recovery underwent a state switch. The lower-tail criterion identifies a distinct subset with evidence of directed departure. The analysis does not identify the destination configuration or the mechanism producing the switch; 'specific' refers to its departure from the arbitrary-restoration expectation. This observation raises the question of whether a high baseline-taxon depletion index can induce a state-switch-like endpoint. Utilizing the community stability landscape framework, we hypothesize that a substantial perturbation directly impacting the community state may enhance the probability of the state transitioning beyond the basin of attraction. To test this hypothesis, we define the 'antibiotic Impact Index' (AII) for a specific subject as follows:

$$\mathrm{AII} = 1 - \frac{|\mathbf{b}^* \cap \mathbf{a}^*|}{|\mathbf{b}^*|} \tag{6}$$

In this context, $\mathbf{a}^*$ and $\mathbf{b}^*$ denote the binary representations of $\mathbf{a}$ (the sample indicative of the antibiotic state) and $\mathbf{b}$ (the sample indicative of the baseline state), respectively. The expression $|\mathbf{b}^* \cap \mathbf{a}^*|$ refers to the number of species common to both the antibiotic and baseline states, whereas $|\mathbf{b}^*|$ indicates the species richness within the baseline state. The Antibiotic Impact Index quantifies the fraction of baseline-detected taxa that are not detected during antibiotic treatment. It is therefore an operational depletion index that depends on sequencing depth, feature resolution, and the applied detection threshold; it should not be interpreted as proof that those taxa were biologically eliminated. Higher AII values indicate greater antibiotic-associated loss from detection among baseline taxa. Subjects meeting the lower-tail switch criterion were concentrated at high AII values in both datasets (Supplementary Fig. 10), linking directed shifts toward specific alternative configurations to severe antibiotic-associated depletion of detectable baseline taxa. Nevertheless, several subjects with similarly high AII values did not meet the switch criterion, indicating that severe detectable-taxon depletion increases the apparent likelihood of switching but is not sufficient to determine the outcome.

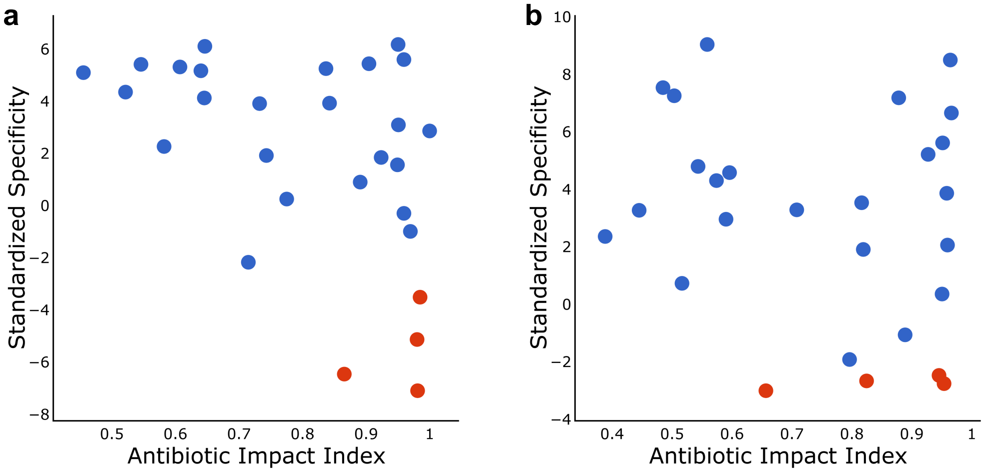


**Supplementary Figure 10: Relationship between the Antibiotic Impact Index and directed switching toward a specific alternative state.** The Antibiotic Impact Index is the fraction of baseline-detected taxa not detected during antibiotic treatment. Red points identify subjects whose own-baseline specificity was significantly below the random-sampling null distribution after multiple-testing correction. Because this null represents arbitrary restoration from the cohort-wide baseline pool, these lower-tail cases indicate a directed, non-random departure toward a specific alternative configuration. Such cases were uncommon and concentrated among subjects with high AII values. **a**, Sewunet et al. [2] cohort. **b**, Messaoudene et al. [3] cohort.

### 11 Cross-species/subject re/establishment analysis for different abundances

In the main text, we show that the cross-species re-establishment empirical probability (returned species) exceeded the cross-species establishment empirical probability (new species) in both analyzed datasets. One possible explanation is that new species generally have lower relative abundances than returned species, and the difference between the probabilities is due to the fact that new species generally have a lower likelihood of being detected. To address this, we repeated the analysis using different minimal relative-abundance thresholds. As shown in Supplementary Fig.11a, increasing the threshold did not change the observed pattern.

In addition, using the species analyzed in Fig. 4**f** of the main text, we tested whether their relative abundance in the second-to-last post-antibiotic sample differed depending on whether they were classified as colonizers or transient. Supplementary Fig.11b shows no significant differences in relative abundance, either for new species or for returned species.

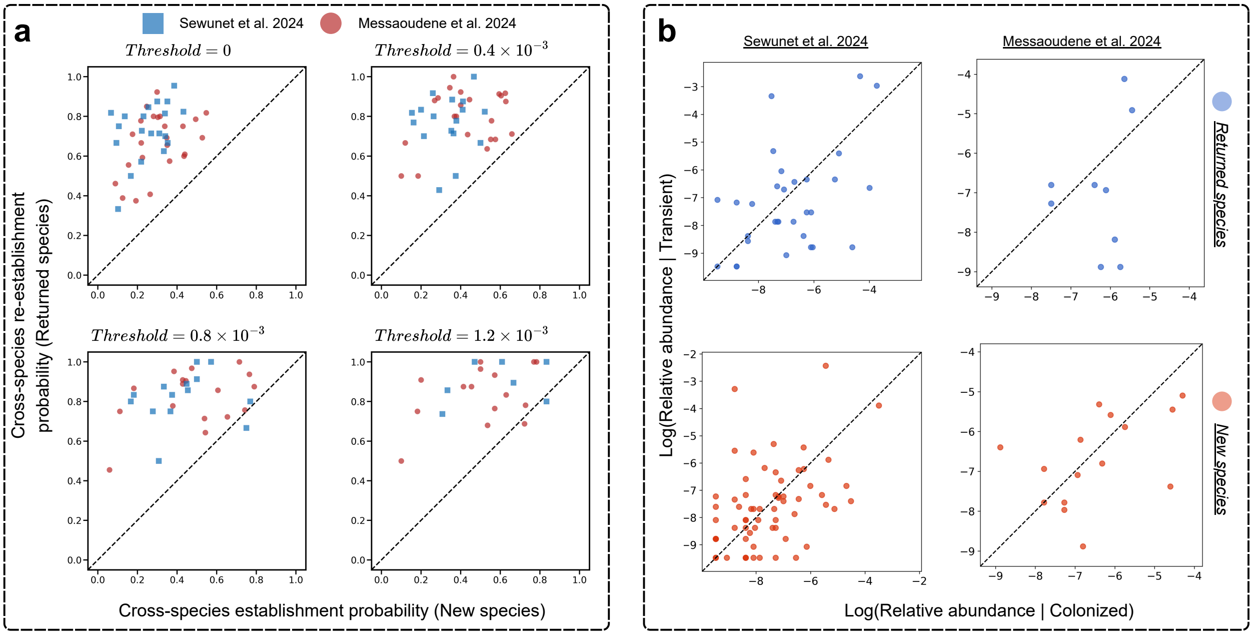


**Supplementary Figure 11:** **a**, The panels illustrate the likelihood of new species and returned species observed in the second-to-last sample during the follow-up period to establish or re-establish, meaning they were detected in the final sample. Different panels present results for varying relative abundance thresholds ($0, 0.4 \cdot 10^{-4}, 0.8 \cdot 10^{-4}, 1.2 \cdot 10^{-4}$), where species with relative abundance below the threshold in the second-to-last sample were excluded. The findings are shown for subjects from Ref. [2] and Ref. [3]. For Ref. [2], the percentages of excluded species at non-zero thresholds are 58% (34%), 74% (50%), and 81% (59%) for new species (returned species). For Ref. [3], the corresponding percentages are 36% (10%), 60% (27%), and 73% (40%) for new species (returned species). Cases where probabilities were estimated based on 5 or fewer species were excluded. **b**, Relative-abundance comparison between colonization and transient outcomes. Each point represents a species classified either as a new species that was observed as both a colonizer and transient in different subjects, or as a returned species first detected in the penultimate follow-up sample and then either persisted or disappeared in the last post-antibiotic sample. For each species, relative abundance was calculated separately for colonization and transient cases. Log-transformed relative abundances are shown, and the dashed line indicates equality between the two outcomes.

#### 12 Historical contingency simulation using a consumer-resource model

To illustrate the historical contingency mechanism using a mechanistic consumer-resource model, we conducted the following simulation, using a similar methodology in parameter generation that is used in Ref [7]. We considered a regional pool of $N$ species competing for $N_R$ resources. The dynamics of species abundances and resource concentrations were described by

$$\frac{dR_\alpha}{dt} = \rho_\alpha - R_\alpha \sum_{i=1}^{N} C_{\alpha i} S_i, \tag{7}$$

and

$$\frac{dS_i}{dt} = S_i \left( \sum_{\alpha=1}^{N_R} G_{\alpha i} R_\alpha - \mu_i \right), \tag{8}$$

where $R_\alpha$ is the concentration of resource $\alpha$, $S_i$ is the abundance of species $i$, $C_{\alpha i}$ is the consumption rate of resource $\alpha$ by species $i$, $G_{\alpha i}$ is the growth rate of species $i$ on resource $\alpha$, $\rho_\alpha$ is the resource supply rate, and $\mu_i$ is the mortality rate of species $i$.

The growth matrix $\mathbf{G} \in \mathbb{R}_{\geq 0}^{N_R \times N}$ was generated by sampling each element independently from $U[0, 1]$. We then generated an independent matrix $\mathbf{C}_0 \in \mathbb{R}_{\geq 0}^{N_R \times N}$, with elements also sampled from $U[0, 1]$, and defined the consumption matrix as

$$\mathbf{C} = \Upsilon \mathbf{G} + \sqrt{1 - \Upsilon^2} \mathbf{C}_0, \tag{9}$$

where $\Upsilon \in [0, 1]$ controls the similarity between growth and consumption. The entries of $\mathbf{C}$ and $\mathbf{G}$ were sampled independently and identically distributed in this way.

Next, we sampled a feasible full-pool fixed point by drawing $R_\alpha^*$ and $S_i^*$ from $U[0.01, 1]$ by sampling each component independently. The mortality vector and resource supply vector were then determined from the fixed-point conditions: $\mu_i = \sum_{\alpha=1}^{N_R} G_{\alpha i} R_\alpha^*$, and $\rho_\alpha = R_\alpha^* \sum_{i=1}^{N} C_{\alpha i} S_i^*$. This construction ensures that $(\mathbf{R}^*, \mathbf{S}^*)$ is a feasible equilibrium of the full species pool.

We then generated an initial community matrix $\mathbf{Y}^{(0)} \in \mathbb{R}_{\geq 0}^{M \times N}$, where each row contained $k$ non-zero species chosen randomly from the regional pool. For selected species, the initial abundance was set to the corresponding value in $\mathbf{S}^*$, while all other species were set to zero. Initial resource concentrations were sampled as small perturbations around $\mathbf{R}^*$: $R_\alpha^{(0)} = R_\alpha^* \xi_\alpha$, where $\xi_\alpha \sim U[0.95, 1.05]$.

We numerically integrated the consumer-resource equations for each initial community and obtained the first steady-state matrix $\mathbf{Y}^{(p)}$. Values smaller than the minimum abundance threshold $\eta$ were set to zero, and each row of $\mathbf{Y}^{(p)}$ was normalized such that $\sum_{i=1}^{N} Y_{ji}^{(p)} = 1 \quad \forall j$. We then selected one row $\mathbf{y}$ from $\mathbf{Y}^{(p)}$ as the test sample. For the test sample and for all other rows in $\mathbf{Y}^{(p)}$, species with zero abundance were reintroduced with a small abundance $\epsilon$, representing immigration from the regional pool. The consumer-resource equations were then integrated again, yielding the post-immigration steady state of the test sample, denoted by $\mathbf{y}^{(s)}$, and the post-immigration cohort matrix $\mathbf{Y}^{(s)}$. Finally, values smaller than $\eta$ were set to zero and the rows of $\mathbf{Y}^{(s)}$ were normalized.

We defined $\{\phi = i \mid y_i > 0 \cap i \mid y_i^{(s)} > 0\}$, as the set of species present both before and after immigration in the test sample, and $\nu = \{i \mid y_i = 0 \cap i \mid y_i^{(s)} > 0\}$, as the set of newly established species. Finally, we applied the *Similarity correlation* analysis, calculating the *sCorr* value using $\mathbf{y}^{(s)}$ as the test sample and the other samples in $\mathbf{Y}^{(s)}$ as references.

In our simulations, we used the parameters $(N, N_R, M, k, \Upsilon, \eta, \epsilon) = (40, 60, 500, 20, 0.35, 10^{-4}, 10^{-10})$. The condition for a steady state was defined as $\max_i \dfrac{|dS_i/dt|}{|S_i| + \varepsilon_0} < \delta$, where $\varepsilon_0 = 10^{-12}$ is a small regularization constant and $\delta = 6 \times 10^{-4}$. If the steady-state condition was not satisfied by $t = 2.5 \times 10^4$, the sample was rejected. Numerical integration was performed using the `solve_ivp` function from the `scipy.integrate` library in Python, with the explicit Runge-Kutta `RK45` method. The results derived from this simulation are presented in Supplementary Fig. 12.

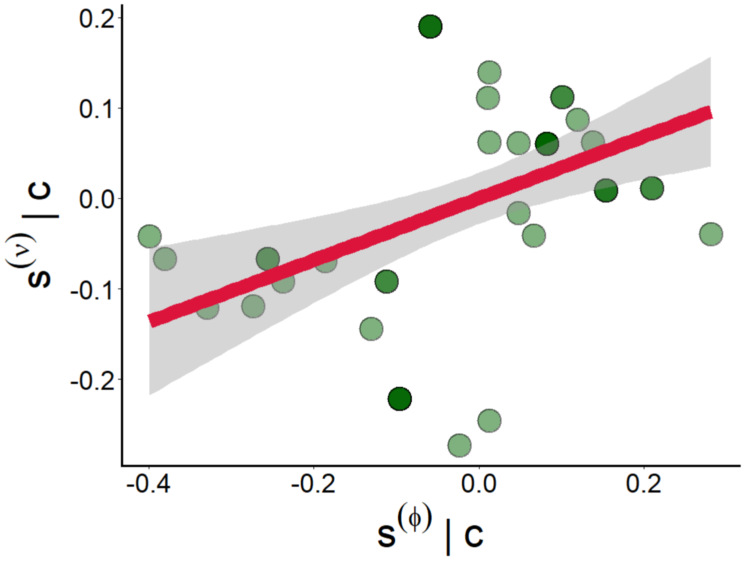


**Supplementary Figure 12:** The outcomes of the similarity correlation conducted through the historical-contingency simulation employing the consumer-resource model ($r = 0.43$, $p < 10^{-3}$).

### 13 Functional association between new colonizers and baseline species in shotgun metagenomic data

In the main text, the functional association between new colonizers and baseline species was assessed using PICRUSt2-based functional profiles inferred from 16S rRNA gene sequencing data. Here, we repeated the analysis using functional profiles derived directly from shotgun metagenomic data of the Yaffe et al. cohort [1] (HUMAnN pathway contributions; see Methods in the main text). The results are consistent with those reported in the main text: colonizers were more functionally similar to the baseline species than transient species, with AUC values of approximately 0.6 for all three reference groups (Supplementary Fig. 13). Unlike in the main text, we present only the pooled analysis across subjects, since the number of transient and colonizing taxa per subject was too low to obtain sufficient statistical power for subject-level tests.

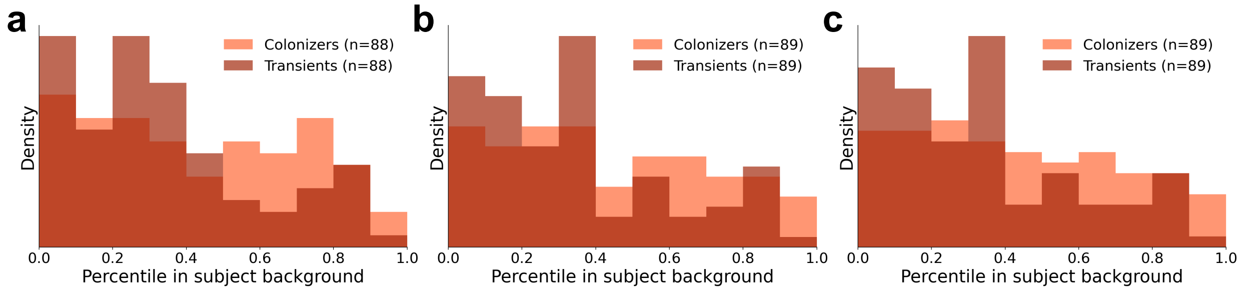


**Supplementary Figure 13: Functional similarity of colonizing and transient species to baseline species in metagenomic data.** Histograms compare colonizers (orange) and transients (dark red) against subject-specific reference groups: **(a)** lost baseline species, **(b)** remaining baseline species, and **(c)** all baseline species. Analyses included 57 antibiotic-treated participants from Yaffe et al. [1]. Baseline species were detected in all four baseline samples; lost species were additionally undetected on day 5 (the final antibiotic-treatment sample) and throughout recovery. Group (b) did not require uninterrupted detection. Colonizers and transients were undetected throughout baseline and antibiotic treatment. Colonizers were detected on both days 28 and 77; transients occurred in one to two recovery samples before day 77 and were undetected on day 77. Binary functional profiles were derived from species-stratified HUMAnN pathway contributions, with positive contributions indicating presence. Community-total, unclassified, and unmapped/unintegrated entries were excluded, and contributors mapping to the same species were combined by pathway union. Species lacking usable profiles were excluded. Reference profiles came from day 0; each colonizer or transient contributed one profile per participant, selected at its highest relative abundance among recovery samples with usable profiles (latest sample for ties). Optimal one-to-one abundance matching without replacement across participants maximized pair counts, then minimized total absolute differences in $\log_{10}$ relative abundance within a 0.2 caliper (approximately 1.58-fold). Matching allowed pairs from different participants and selected 89 pairs from 119 colonizer and 121 transient observations. Each species' score was its unweighted mean Jaccard similarity (shared pathways divided by their union) to its participant's reference species. Scores were converted to participant- and reference-specific background percentiles. Background profiles represented species detected on day 77 in other participants, selected using the same highest-abundance recovery rule, excluding the focal participant's baseline, colonizer, and transient species. Background species were scored against the focal participant's reference group; percentiles equalled the fraction of background scores below the focal score plus half the fraction tied. Higher percentiles therefore indicate greater relative similarity to the reference group. Comparisons required at least one usable reference species and 30 background profiles. Pooled percentiles are shown in ten equal-width bins over $[0, 1]$, with unit-area normalization per group. Tests required at least eight observations per pooled group and retained only pairs with both members scored. AUC measures the probability that a colonizer's percentile exceeds a transient's, counting ties as one-half. Two-sided permutation $p$ values used 10,000 within-pair label swaps and deviations from the mean permutation AUC. Results: **(a)** 88 pairs, AUC = 0.590 (95% CI, 0.509-0.671), $p = 0.018$; **(b)** 89 pairs, AUC = 0.600 (95% CI, 0.525-0.677), $p = 0.011$; **(c)** 89 pairs, AUC = 0.603 (95% CI, 0.528-0.680), $p = 0.008$.

### 14 Prediction of individual-specific recovery using regularized linear regression

We investigated whether the microbial community observed during antibiotic treatment could predict the degree of individual-specific microbiome recovery. The predictor matrix, denoted by $\mathbf{X} \in \mathbb{R}^{M \times N}$, contained the microbial abundance profiles of $M$ subjects at baseline, before antibiotic treatment, where rows represented subjects and columns represented microbial taxa. The regression target was a standardized Jaccard similarity (see the main text, Fig. 2, and the Methods section), a measure of individual-specific recovery.

**Leakage-aware leave-one-out cross-validation**. Predictive performance was evaluated using leave-one-out cross-validation (LOOCV). In each outer fold, one subject was held out as the validation sample and the remaining $M - 1$ subjects were used for model training. Because the target, the standardized Jaccard similarity, depends on similarities to the baseline reference cohort, the target was recalculated separately inside every outer cross-validation fold. This prevented the baseline of the held-out subject from contributing to the target values used for training.

For the held-out subject, the target was calculated relative to the same allowed reference set. Since the held-out subject's own baseline was already omitted from the corresponding row of the comparison matrix, all remaining allowed reference baselines were used. Additional independent baseline samples that were not represented in $\mathbf{X}$ remained available in every cross-validation fold.

**Data preprocessing**. All preprocessing operations were incorporated into a `Scikit-learn` pipeline and were fitted using only the training samples in each cross-validation fold. First, taxa were filtered according to their prevalence. A taxon was retained if it had nonzero abundance in at least a minimum fraction of the training samples; this prevalence threshold was treated as a hyperparameter and tuned by cross-validation (see below). Next, the retained microbial abundances were transformed using the centered log-ratio transformation. A pseudocount $\delta = 10^{-6}$ was first added to each abundance value. For subject $i$, the transformed abundance of taxon $j$ was $\mathrm{clr}(\mathbf{X}_{ij}) = \log(\mathbf{X}_{ij} + \delta) - \frac{1}{P_i} \sum_{k=1}^{P_i} \log(\mathbf{X}_{ik} + \delta)$, where $P_i$ is the number of retained taxa. Finally, every retained CLR-transformed feature was standardized using the mean and standard deviation estimated from the training samples. The fitted prevalence filter, CLR transformation, and standardization parameters were then applied unchanged to the held-out subject.

**LASSO regression and hyperparameter optimization**. The transformed microbial profiles were used to fit a LASSO regression model:

$$\widehat{\boldsymbol{\beta}} = \arg\min_{\boldsymbol{\beta}} \left[ \frac{1}{2M_{\text{train}}} \sum_{i=1}^{M_{\text{train}}} \left( z_i - \beta_0 - \mathbf{x}_i^{\mathsf{T}} \boldsymbol{\beta} \right)^2 + \alpha \sum_{j=1}^{N_{\text{train}}} |\beta_j| \right]. \tag{10}$$

The $L_1$ penalty shrinks regression coefficients and can set some coefficients exactly to zero, thereby performing embedded feature selection.

Hyperparameters were optimized separately inside every outer LOOCV training set using an inner shuffled $K$-fold cross-validation procedure. By default, five inner folds were used. A grid search evaluated all combinations of two hyperparameters: the LASSO regularization strength $\alpha \in \left\{10^{-4}, \ldots, 10^{0}\right\}$, taken over 200 logarithmically spaced values, and the prevalence-filter threshold applied to each microbial feature, taken over a small grid of candidate values. The hyperparameter combination minimizing the inner cross-validated mean squared error was selected. The complete preprocessing pipeline and LASSO model were then refitted on all subjects in the outer training set and used to predict the standardized Jaccard similarity of the held-out subject.

**Performance evaluation**. After repeating this procedure for every subject, each subject had one prediction obtained from a model that had not been trained on that subject. Predictive performance was evaluated using the pooled out-of-fold predictions. The following metrics were calculated:

the root mean squared error,

$$\text{RMSE} = \sqrt{\frac{1}{M} \sum_{i=1}^{M} (z_i - \widehat{z}_i)^2}. \tag{11}$$

and the mean absolute error,

$$\text{MAE} = \frac{1}{M} \sum_{i=1}^{M} |z_i - \widehat{z}_i|, \tag{12}$$

Pearson correlation was used to quantify the linear association between observed and predicted scores. The results obtained for subjects from Refs. [2, 3] are shown in Supplementary Fig. 14

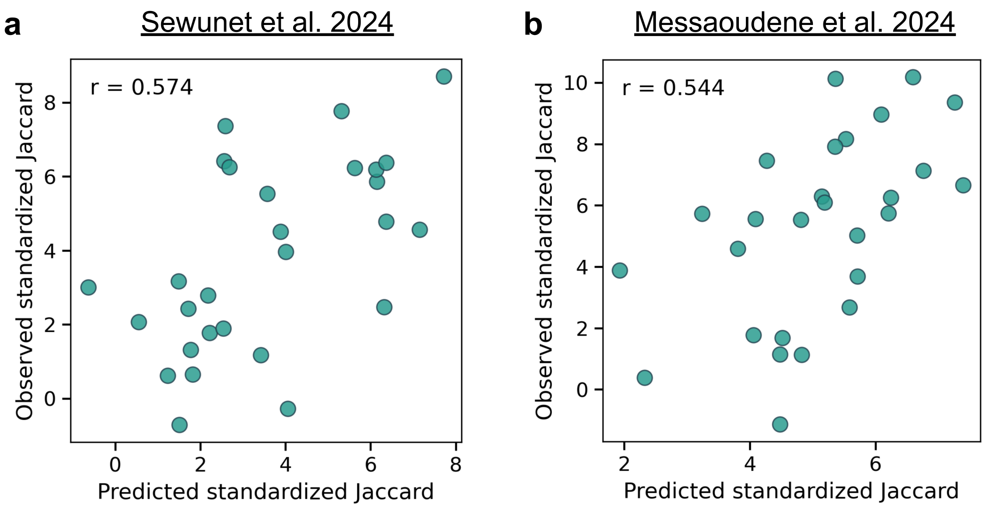


**Supplementary Figure 14:** Observed versus predicted standardized Jaccard similarity scores obtained via leave-one-out cross-validation. **a**, Results for subjects from Ref. [2] using taxonomic profiles as features (Pearson correlation: 0.574, RMSE: 2.201, MAE: 1.697). **b**, Corresponding metrics for subjects from Ref. [3] using taxonomic profiles (0.544, 2.543, and 2.151, respectively).

[2] Tsegaye Sewunet, Mohammad Razavi, Staffan Rosenborg, Angela Camporeale, Michael Nowak, David Melnick, Leanne B Gasink, Paul B. Eckburg, Ian A Critchley, Carl Erik Nord, and Christian G. Giske. Effect of tebipenem pivoxil hydrobromide on the normal gut microbiota of a healthy adult population in sweden: a randomised controlled trial. *The Lancet. Microbe*, 2024.
